# Targeting dual signaling pathways in concert with immune checkpoints for the treatment of pancreatic cancer

**DOI:** 10.1101/860247

**Authors:** Erik S Knudsen, Vishnu Kumarasamy, Sejin Chung, Paris Vail, Stephanie Tzetzo, Amanda Ruiz, Mukund Seshadri, Scott I Abrams, Jianmin Wang, Agnieszka K Witkiewicz

**Affiliations:** Center for Personalized Medicine, Roswell Park Comprehensive Cancer Center, Buffalo NY; Molecular & Cellular Biology, Roswell Park Comprehensive Cancer Center, Buffalo NY; Biostatistics and Bioinformatics, Roswell Park Comprehensive Cancer Center, Buffalo NY; Immunology, Roswell Park Comprehensive Cancer Center, Buffalo NY; Cancer Center, University of Arizona, Tucson AZ; Oral Oncology, Roswell Park Comprehensive Cancer Center, Buffalo NY; Pathology, Roswell Park Comprehensive Cancer Center, Buffalo NY

## Abstract

Pancreatic cancer harbors a poor prognosis due to the lack of effective systemic therapies. Here we interrogated means to target key effector pathways down-stream from KRAS. We found that combination treatment with MEK and CDK4/6 inhibitors was effective across a broad range of PDX models in delaying tumor progression. These effects were associated with stable cell cycle arrest, as well as the induction of multiple genes associated with interferon response and antigen presentation in an RB-dependent fashion. Using single cell sequencing and complementary approaches, we found that the combination of CDK4/6 and MEK inhibition had a significant impact on increasing T-cell infiltration and altering myeloid populations, while potently cooperating with immune checkpoint inhibitors. Together, these data indicate that there are canonical and non-canonical features of CDK4/6 and MEK inhibition that impact on the tumor and host that can contribute to durable control for tumors in combination with immune checkpoint inhibitor therapy.

## INTRODUCTION

Pancreatic ductal adenocarcinoma (PDAC) has a universally poor prognosis that has proven resistant to multiple targeted interventions^1, 2^. KRAS mutations dominate the genetic landscape of pancreatic cancer and exist in concert with a number of high-potency genetic events (e.g. CDKN2A loss, TP53 mutation, SMAD4 loss) that contribute to the aggressive nature of PDAC^3-5^. Clinical trials of agents targeting RAS pathway in pancreatic cancer via MEK or other effector pathway inhibitors have largely failed to demonstrate^6, 7^. Additionally, even genetic strategies to ablate KRAS in preclinical models of pancreatic cancer only lead to transient remission^8^. Thus, targeting multiple pathways to control oncogenic signaling is likely important for limiting adaptation around inhibiting critical genetic nodes in PDAC.

Many oncogenes, including KRAS impinge on the cell cycle through the activation of CDK4 and CDK6 kinases^9-11^. In PDAC, deletion or mutation of the CDKN2A gene leads to the loss of p16ink4a that is an endogenous CDK4/6 inhibitor. In the context of KRAS-driven tumorigenesis, loss of p16ink4a is critical for the bypassing oncogene-induced senescence^12^ and thus effective suppression of CDK4/6 activity could be particularly relevant in the context of pancreatic cancer^13^. Interestingly, in preclinical studies PDAC models are surprisingly resistant to CDK4/6 inhibition^14, 15^. However, combination with MTOR or MEK inhibitors have been reported to enhance response to CDK4/6 inhibition in PDAC and other RAS-driven tumor types^15-17^. In spite of the potency of such combinations in enforcing cell cycle withdrawal, tumor cells continue to survive and as a result acquired resistance represents a significant challenge with such cytostatic combinations.

Immunotherapy has emerged as an important treatment modality in a number of tumor types including those harboring RAS mutation (e.g. lung cancer and melanoma)^18, 19^. However, single agent immunotherapy treatments have generally not been successful in pancreatic cancer^20-22^. This lack of response may in part be attributed to the presence of dense desmoplastic stroma, the relatively low level of quality neo-epitopes, and the activation of multiple immune suppressive features of the tumor microenvironment^21-25^. Understanding the complex interplay between tumor cell and stromal compartment and agents that target these components have been utilized in combination with immune modulating treatments and have shown promising results^26, 27^. Rationally combining targeted and immune-modulatory agents for the treatment of PDAC could represent an important advance in yielding long-term disease control for pancreatic cancer patients.

## METHODS

### Cell culture

Pancreatic ductal adenocarcinoma cells were grown in Keratinocyte SFM with 0.2 ng/mL EFG, 30 *µ*g/mL bovine pituitary extract (Life Technologies, 10744019), and 2% fetal bovine serum on collagen coated (Millipore 08-115), tissue culture treated plates as published^24, 28^. The 4662 model was provided by Dr. Robert Vonderheide, and cultured in DMEM+10%FBS as reported^29, 30^.

### Cell proliferation assay

Cell proliferation following different treatments was determined using chemiluminescent BrdU ELISA kit (Roche, catalog # 11669915001) as described by the manufacturer. Luminescence was read on a Biotek Synergy 2 plate reader.

### Immunoblot analysis

Primary antibodies for immunoblot analysis were purchased from Cell Signaling Technology include p-RB (S807/S811) (8516S), RB (9313S) (p-Akt (S473) (4070S), Akt (4691), pS6 (S235/236) (2211), S6 (2217), cyclin E1 (4129S), p27 (2552S). Anti-Actin (SC-47778), anti-pERK (Y204) (SC-7383), anti-ERK (SC-514302), anti-cyclin D1 (SC-20044) and anti-cyclin A (SC-271682) were purchased from Santa Cruz Biotechnology. The whole-cell extracts were prepared by lysing the cells with RIPA lysis buffer (Santa Cruz Biotechnology, SC-24948A) in the presence of 1X Halt protease inhibitor (Thermo Fisher) and 1 mM PMSF (Sigma). The extracted proteins (20 *µ*g) were resolved by SDS-PAGE and transferred to PVDF membranes, which were then incubated with primary antibodies at 4°C overnight, followed by incubation with HRP tagged anti-mouse or anti-rabbit secondary antibodies at room temperature up to 1 hour. An enhanced chemiluminescence kit (Thermo Fisher, 34076) was used to detect the immuno-reactive bands.

### CDK2 kinase assay

To analyze the CDK2 kinase activity, primary PDAC cells were lysed using the kinase lysis buffer (50 mM HEPES-KOH pH 7.5, 150 mM NaCl, 1 mM EDTA, 1mM DTT, 0.1% Tween-20) in the presence of 1X Halt protease inhibitor (Thermo Fisher) and 1 mM PMSF (Sigma). Active CDK2 complex was immunoprecipitated by incubating 300 *µ*g of the lysate with 5 *µ*g of anti-CDK2 (Santa Cruz Biotechnology; SC-6248) overnight at 4°C. Normal mouse IgG1 (Cell Signaling Technology, 5415) was used as a control. Protein G agarose-beads were added to each IP samples and incubated up to 4 hour at 4°C. Protein immunocomplexes were washed 3 times with the kinase lysis buffer and 2 times with kinase reaction buffer (40 mM Tris-HCl pH 8, 20 mM MgCl_2_, 0.1 mg/mL BSA, 50 *µ*M DTT). Kinase reactions were carried out in 100 *µ*l of kinase buffer in the presence of 100 *µ*M ATP and 0.5 *µ*g of RB protein as substrate by gently shaking at room temperature up to 30 minutes. The resulting phosphorylated RB protein was detected by western blotting using anti-p-Rb (S807/S811) antibody (Cell Signaling Technology, 8516S).

### FUCCI (fluorescent, ubiquitination-based cell cycle indicator) Transfection

ES-FUCCI, a gift from Pierre Neveu (Addgene plasmid #62451), was transfected to 226 cell line using Lipofectamine 3000 Reagent (Invitrogen LSL3000001) by manufacturer’s protocol. Transfected cells were selected using Hygromycin and sorted with FACSAria.

### Drug Screen

IncuCyte S3 Live-Cell Analysis System (Essen Biosciences) was utilized for live cell analysis. Primary PDAC cells were labeled with H2B-GFP and seeded in 384-well collagen-coated plates. Cells were treated with DMSO or Palbociclib (100nM) for 24 hours prior to library screen of over 300 cancer drugs at 100nM. Two phase and fluorescent images per well were captured hourly for 72 hours at 10X magnification. Essen Bioscience software was used to quantify number of cells per well and normalized to DMSO treated cell proliferation. Data was exported to Prism 7 (GraphPad) for statistical analyses.

### Gene Expression Analysis

RNA sequencing read counts were normalized using edgeR package in R. The log fold change and Student’s two-sided t test p-value were calculated (assuming equal variance) for each treatment relative to the control from the normalized reads across different cell line and PDX model. Volcano plots used cutoffs for log fold change as specified in the figure legends. Genes that are upregulated and downregulated in the palbociclib+tramentinib condition were used for gene ontology analysis using ENRICHR, and gene set enrichment analysis was performed as previously published. For the analysis of TCGA data expressed genes in palbociclib+tramentinib related to immune function and cell cycle were applied to the pancreatic cohort and clustered based on k-means. Kaplan-Meier analysis for overall survival comparing the clusters was performed using the survival package in R.

### Immunohistochemical, multi-spectral imaging

Immunohistochemistry on mouse tissues was performed using standard procedures with antibodies against: CD8 (ab209775-1:100), CD163 (ab182422-1:250), Ki67 (RM-9106S1-1:150). Staining was performed on a Leica auto-stainer. Images and quantification were performed using Aperio (Leica Biosystems). Multispectral panels including the above antibodies were developed and images captured on a Vectra Polaris Instrument (Perkin Elmer). Phenotype counts were determined using InFORM software (Perkin Elmer).

### Mice and Patient-derived xenografts

NSG (Jackson Laboratories) mice were maintained in the University of Arizona animal care facility. Mice were both male and female and used between 10 and 20 weeks of age for engraftment. All animal care, treatment, and sacrifice were approved by the University of Arizona Institutional Animal Care and Use Committee (IACUC) in accordance with the National Institutes of Health (NIH) Guide for the Care and Use of Laboratory Animals. Mice were implanted subcutaneously with early passage PDX tumor fragments that have never been placed into culture. When tumors reached a volume of ∼200 mm^3^ they were randomized to treatment cohorts such that the starting volume across the cohorts was similar. Mice were treated for 3 weeks by gastric gavage with single agent or a combination. Palbociclib (PD-0332991, 100 mg/kg) diluted in 50 mM lactate buffer at pH 4.0, Trametinib (0.5 mg/kg) diluted in 0.5% hydroxypropyl cellulose and 0.2% Tween 80. Mice were treated daily for 5 days, followed by a 2-day break during 1^st^ week, and then every other day during 2^nd^ and 3^rd^ weeks for a total treatment time of 3 weeks. Tumor size was assessed every 2 days using digital calipers. Mice were sacrificed at either 6 days of treatment, 21 days of treatment, or 2 weeks after the end of treatment. Tumor measures were carried out independently by multiple laboratory members.

### Syngeneic tumor model

C57BL/6J mice were subcutaneously injected with 4662 cells (3 × 10^6^). When tumor volumes reached 150mm^3^ mice were randomized to control, Trametinib, Palbociclib, and combination. Other groups were anti-PDL1, Palb +Tram + IgG, and Palb + Tram + anti-PDL1. Tumors were measured every other day with caliper and volume was calculated with the following equation: V = 0.5 ([greatest diameter] × [shortest diameter]^2^). Control group was given vehicle, Palbociclib was prepared in 50mM Na-Lactate Buffer (pH=4), Trametinib was prepared in 0.5% hydroxypropyl Cellulose and 0.2% Tween 80 in water, and dilution buffer was used for anti-PDL1 as well as IgG. Palbociclib and Trametinib were administered by gastric gavage. Anti-PDL1 and IgG were administered by intraperitoneal injection. Body weight was measured every other day. All the treatments lasted for 21 days or until tumor volumes reached 2000mm3. In mice where the tumor completely regressed, treatment was ceased, and the mice were monitored up to 30 days for tumor outgrowth. At the end of 30^th^ day, mice were re-challenged with 3×10^5 4662 cells and the tumor growth were monitored up to 45 days. Orthotopic models were performed as previously published^31^, mice were randomized at ∼100 mm^3^ as determined by MRI imaging^32^.

### Single cell sequencing and analysis

C57BL6/J mice harboring 4662 tumors were sacrificed on treatment and dissociated by mincing and digestions with liberase (Sigma # 541020001) (2×15’). Resultant cells were stained with anti-mouse CD45 BUV395-conjugated antibody (BD Biosciences # 564616) and viability dye eFluor780 (Thermo Fisher #65-0865-14) and subjected to sorting. Single cell libraries are generated using the 10X Genomics platform. Cell suspensions are first assessed with Trypan Blue using a Countess FL automated cell counter (ThermoFisher), to determine concentration, viability and the absence of clumps and debris that could interfere with single cell capture. Cells are loaded into the Chromium Controller (10X Genomics) where they are partitioned into nanoliter-scale Gel Beads-in-emulsion with a single barcode per cell. Reverse transcription is performed and the resulting cDNA is amplified. The full-length amplified cDNA is used to generate gene expression libraries by enzymatic fragmentation, end-repair, a-tailing, adapter ligation, and PCR to add Illumina compatible sequencing adapters. The resulting libraries are evaluated on D1000 screentape using a TapeStation 4200 (Agilent Technologies), and quantitated using Kapa Biosystems qPCR quantitation kit for Illumina. They are then pooled, denatured, and diluted to 300pM with 1% PhiX control library added. The resulting pool is then loaded into the appropriate NovaSeq Reagent cartridge and sequenced on a NovaSeq6000 following the manufacturer’s recommended protocol (Illumina Inc.).

The raw sequencing data were processed using Cell Ranger software with mouse mm10 reference genome^33^. The filtered gene-barcode matrices which contain barcodes with the Unique Molecular Identifier (UMI) counts that passed the cell detection algorithm were used for Seurat single cell data analysis R package ^*34*^. Cells with low RNA features (< 200) or high RNA feature (>5000) or high mitochondrial RNA contents (10%) were filtered out from the analysis. Data from all samples are merged together and normalized using the SCTransform. Dimension reductions (PCA and UMAP) using the highly variable genes were calculated for clustering analysis using the first 30 principal components. SingleR package was utilized to identify the cell types and RNA velocity analysis were carried out using velocyto R package ^35, 36^.

## RESULTS

In spite of genetic features that would be expected to yield sensitivity to CDK4/6 inhibitors pancreatic cancer cell lines are surprisingly resistant to this treatment^16, 37^. To systematically define cooperating agents, we carried out live cell imaging-based drug screens where the suppression of proliferation is directly monitored by evaluating cell number (Fig 1A and S1). In this screen, MEK inhibitors were highly enriched for cooperating with the CDK4/6 inhibitor palbociclib. Validation studies with the MEK inhibitor trametinib demonstrated that these effects are due to cooperation relative to cell cycle inhibition as determined by BrdU incorporation (Fig 1B). Furthermore, dose response analysis showed that the drug interaction is synergistic as determined by Bliss analysis in multiple patient-derived cell models (Fig 1C and S1). To interrogate the mechanisms related to response, initially we evaluated how CDK4/6 inhibition may impact on features of signaling downstream from KRAS (Fig S1). These data showed that CDK4/6 inhibition had no effect related to canonical signaling through ERK, AKT, or MTOR pathways as determined by phosphorylation of ERK, AKT and S6 respectively (Fig S1). However, the treatment with trametinib resulted in expected suppression of ERK activity and in several models also suppressed activity through the MTOR and AKT pathways (Fig S1), as has been recently reported in other RAS-driven tumors^17^. To determine the features of therapeutic cooperation, we interrogated canonical determinants of cell cycle control. Treatment with CDK4/6 inhibitors lead to the adaptive upregulation of cyclin D1 and cyclin E in the pancreatic cancer cell lines (Fig 1D), consistent with prior findings^37, 38^. These adaptive features of CDK4/6 inhibition were ameliorated with the combined treatment with trametinib that yielded potent blockade of RB phosphorylation and suppression of cyclin A expression (Fig 1D). Flow cytometry analysis using a FUCCI reporter for APC/CDH1 and SCF/SKP2 activity was used to determine the action of drug treatment on cell cycle regulatory activities. These data showed that the combination of CDK4/6 and MEK inhibition yielded a potent suppression of SCF/SKP2, yet APC/CDH1 remained active (Fig 1E and S2). In parallel with these findings, we found that MEK inhibition induced p27Kip1 (Fig S2), which is an important determinant of response to CDK4/6 inhibition^10^. While CDK4/6 inhibition had a modest effect on CDK2 activity, MEK inhibition cooperated with CDK4/6 inhibition to decrease CDK2 kinase activity (Fig 1F and Fig S2). Consistent with the effects on CDK2 and RB activity, gene expression analysis illustrated pronounced suppression of E2F-target genes in multiple cell models with the combination treatment (Fig 1G). Thus, these data support the impact of coordinately targeting both a KRAS effector pathway (MEK1/2) and cell cycle (CDK4/6) to elicit pronounced cell cycle exit.

**Figure 1.**
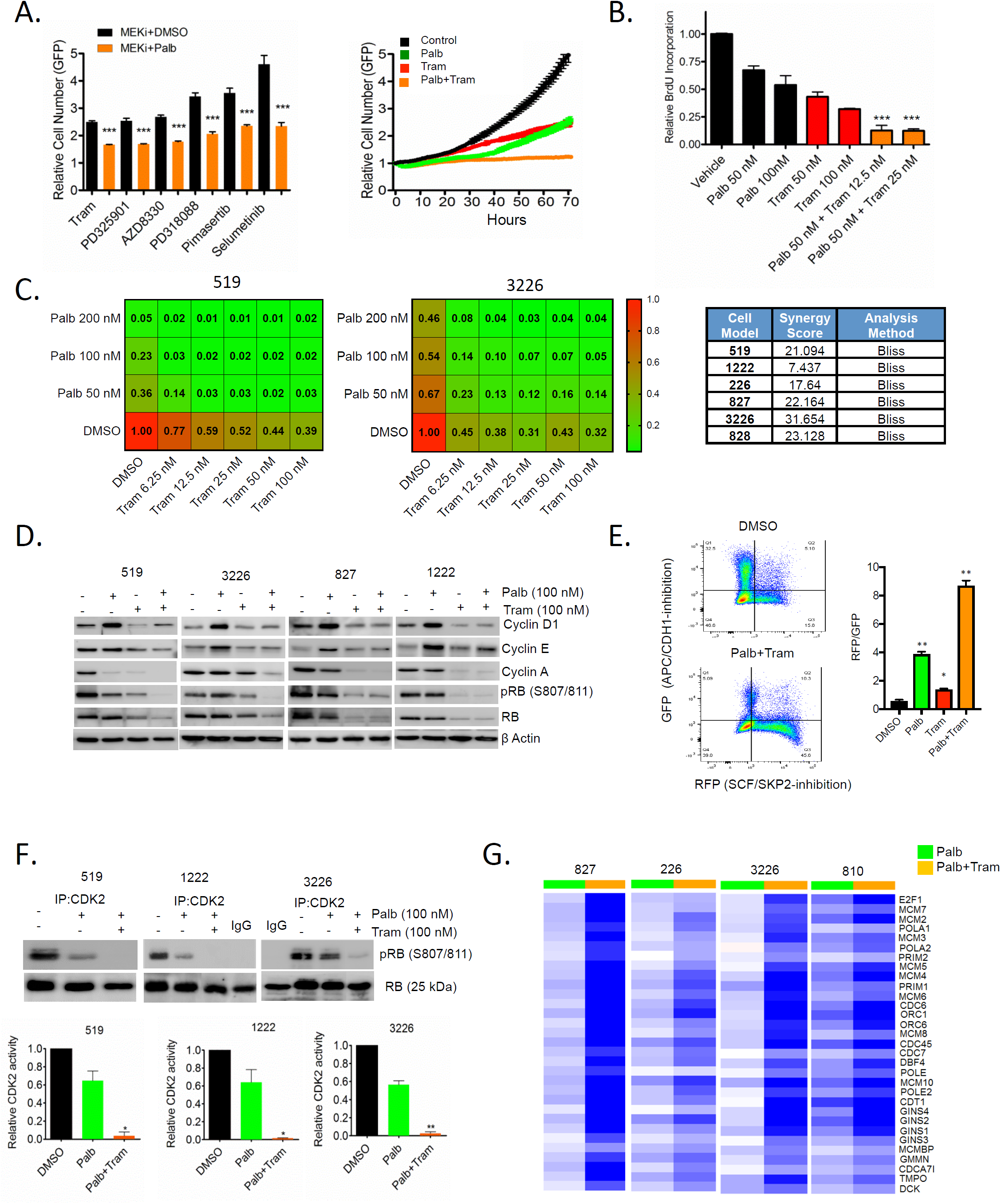
Cooperation between MEK and CDK4/6 inhibition in PDAC cell model: **A.** The 3226 pancreatic cancer cell line, expressing H2B-GFP was treated with palbociclib (100 nM) +/- the indicated MEK inhibitors (250 nM) and relative proliferation was determined by live cell imaging. **B.** Relative BrdU incorporation was determined following 48-hour exposure to palbociclib +/- trametinib at the indicated concentrations in the 3226 cell line. **C.** Isobologram analysis in relatively CDK4/6 inhibitor sensitive (519) and resistiant (3226) cell showing dose-response relationship between palbociclib and trametinib concentrations. Heatmaps represent relative BrdU incorporation after 48 hours treatment. **D.** Immunoblot analysis of the indicated cell-cycle proteins in 519, 3226, 827 and 1222 cell lines that were treated with palbociclib (100 nM) +/- trametinib (100 nM) for 48 hours. **E.** 226 cells were labeled with FUCCI and treated with DMSO, Palbociclib (200nM), Trametinib (50nM), and combination for 48 hours prior to flow cytometry analysis. Cells were counter stained with DAPI for DNA content. Panel shows GFP (APC/CDH1-inhibition) on Y-axis and RFP (SCF/SKP2-inhibition) on x-axis. Graph shows quantitation of RFP to GFP ratio of indicated treatments. * p-value <0.05, ** p-value <0.01 t-test. **F.** *In vitro* CDK2 kinase assays were performed in 519, 1222 and 3226 cell lines following the treatment with palbociclib (100 nM) +/- trametinib (100 nM). Specific kinase activity was determined based on the phosphorylation of an exogenous RB substrate at S807/811 and the band intensities were quantified. Representative blot images and mean and SD are shown (*p<0.5, **p<0.01, as determined by t test). **G.** Heatmap showing the Log-fold change of the indicated cell cycle regulatory genes from cells treated with palbociclib alone or the combination with trametinib.

We utilized a panel of pancreatic cancer PDX models to delineate *in vivo* response to CDK4/6 and MEK inhibition. In multiple models, treatment with single agent MEK, or CDK4/6 inhibition had a transient impact on tumor growth (Fig 2A and 2B). However, the combination significantly reduced tumor growth. Across a large number of individual PDX models (n=10 different PDXs and 368 individual tumors), we found that the combination significantly increased progression-free survival relative to CDK4/6 inhibition alone (Fig 2C). However, with the cessation of therapy at 21 days tumors progressed indicating the reversible cytostatic nature of the treatment. These responses were associated with enhanced suppression of Ki67 proliferation marker by immunohistochemistry, and the repression of E2F-target genes by RNA sequencing (Fig 2D and E). Notably, in the PDX models (as in the cell models) the combination treatment limited phosphorylation of RB, and the expression of cyclin D1 and E (Fig S3). Together, this work shows that the combination with CDK4/6 and MEK inhibition provides a putative therapeutic opportunity for the treatment of PDAC that is associated with profound cell cycle exit.

**Figure 2.**
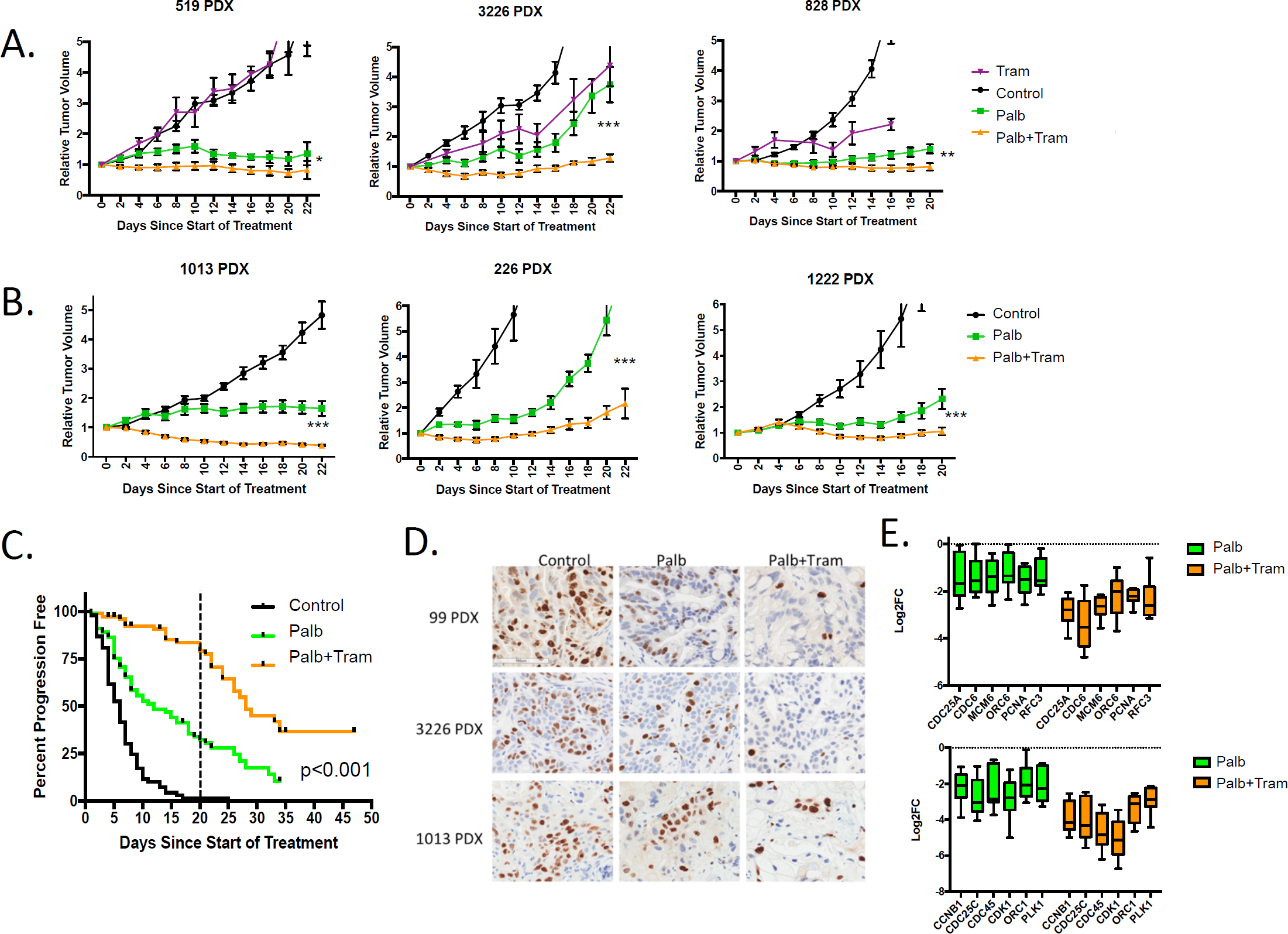
Cooperation between MEK and CDK4/6 inhibition in PDX models of pancreatic cancer: **A.** The indicated PDX models were treated with vehicle, palbociclib or trametinib and the combination. The tumor volume was measured by caliper and the relative tumor volume for the cohorts is plotted as a function of time (*p<0.05, **p<0.01, ***p<0.001). **B.** Additional PDX models were treated with vehicle, palbociclib or palbociclb+trametinib combination. The relative tumor volume for the cohorts is plotted as a function of time (*p<0.05, **p<0.01, ***p<0.001). **C.** Progression-free survival (where progression represents a 50% change in tumor volume) is plotted. Treatment was for 21 days (marked by dashed line), and subsets of animals were followed with the cessation of therapy. **D.** Immunohistochemical staining for Ki67 in three different PDX models treated with palbociclib or the combination with trametinib. **D.** Expression of the indicated cell cycle regulatory genes is shown from the panel of treated PDX models. Log2FC is graphed for each of the indicated genes involved in DNA replication (top panel) or mitosis (bottom panel).

In order to understand means to expand on the cytostatic efficacy of the CDK4/6 and MEK inhibitor combination, we evaluated RNA sequencing data from treated PDAC models. In cell culture models we found that the combination of CDK4/6 and MEK inhibition elicited both the consistent suppression of genes, but also induced an equivalent number of genes (Fig 3A). While the suppressed genes were strongly associated with cell cycle, the upregulated genes were associated with antigen presentation and features of interferon signaling (Fig 3A, 3B and S4). This signature of immune response is similar to that observed either with single agent MEK or CDK4/6 inhibition in other models^26, 39, 40^. In particular, we observed that the combination induced the expression of multiple MHC genes (e.g. HLA-A, HLA-C) and genes involved in interferon signaling (e.g. STAT2 and IRF9) (Fig 3C and S3). The induction of immune-related proteins was dependent on the presence of RB, as the 7310 cell line which is RB-deficient^41^ failed to elicit this response (Fig 3D and S4). Further, evaluating independent treatments illustrated that MEK inhibition suppresses canonical target genes (e.g. DUSP5 and ETV1), as well as cell cycle (e.g. CCNA2 and PLK1), while eliciting the induction of the genes associated with interferon response and antigen presentation (Fig 3E). In this context, the suppression of MEK genes is RB-independent, while cell cycle and induced genes require RB as noted by the cell line 7310 (Fig 3E). The observed changes in gene expression were observed to induce secretion of CCL5 and CXCL10 that are downstream from interferon responses and associated with enhanced T-cell infiltration (Fig S5). The gene expression findings were also recapitulated in PDX models (Fig 3F and S5), where tumor selective transcripts were evaluated through the use of a hybrid mouse/human genome ^42^. The interferon-like response has been suggested to represent a senescence-associated secretory phenotype (SASP). However, the lack no induction of IL6, IL8, and IL1ß in our data and little positive enrichment for the classical SASP signature (Fig S6). In addition to MEK inhibition, MTOR inhibitors can cooperate with CDK4/6 inhibitors in driving cell cycle exit^37^. The TORC1/2 inhibitor TAK228 results in potent cooperation with CDK4/6 inhibition in the suppression of cell cycle-regulated genes in PDX models and cell lines (Fig S7). However, the combination of palbociclib and TAK228 did not recapitulate the induction of the antigen-presentation genes (Fig S7). These data validate that the combination of CDK4/6 and MEK inhibition is distinct in mediating both profound cell cycle exit and immunological response features in pancreatic cancer models. It has been reported in several tumor types that there is an inverse relationship between cell cycle and interferon-related gene expression signatures^43^. Using TCGA data we found that those tumors with high-index for proliferation, but low for immune response are associated with poor prognosis (Fig 3G, 3H and S8). Thus, in principle, shifting the transcriptional program toward that induced by MEK and CDK4/6 inhibition would be associated with improved survival coupling dual biological effects.

**Figure 3.**
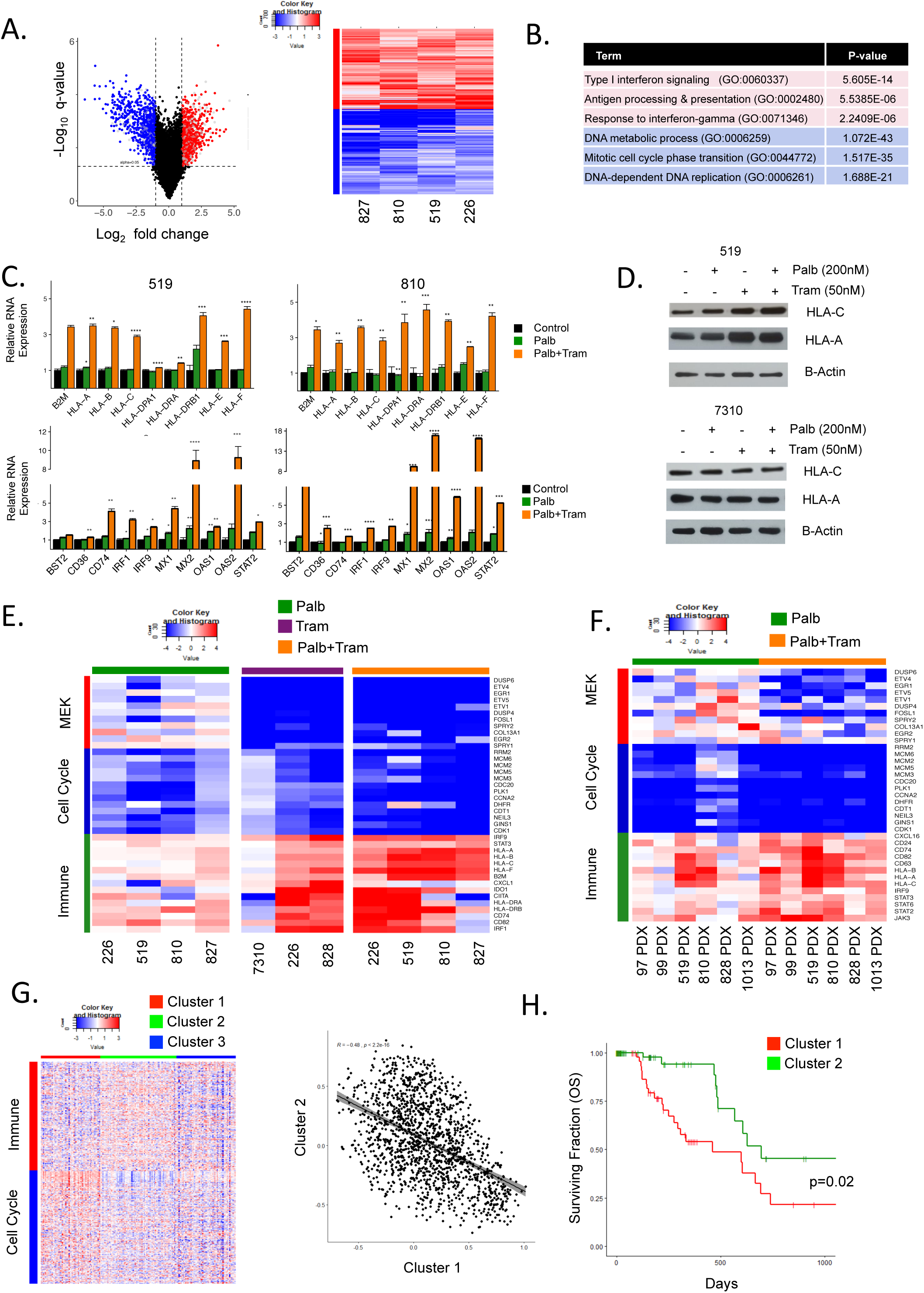
Induction of immunological genes in an RB-dependent manner in PDAC models treated with MEK and CDK4/6 inhibitors. **A.** Common genes expressed in the palbociclib and trametinib treated conditions were determined in cell lines. (left panel) Genes with an average log 2 fold-change greater than 1 and FDR less than 5% were considered to be differentially expressed. The blue symbols denote repressed cell cycle genes and the red symbols denote induced genes associated with the immune system. (right panel) Heatmap comparing the log fold change of genes related to immune function (red-color bar) and cell cycle (blue-color bar) in palbociclib+trametinib treated cell PDAC cell line models. **B.** Top gene ontology terms related to the immune system and cell cycle ranked based on p-value. **C.** Relative mRNA expression of palbociclib (green) and palbociclib+trametinib treated cells for a subset of genes related to antigen presentation and Interferon signaling (*:p<=0.05, **:p<=0.01,***:p<=0.001,****:p<=0.0001). **D.** Immunoblots for the indicated proteins were performed in the 519 cell line, and the RB-deficient 7310 cell line. **E.** Heatmap showing the log fold change of representative MEK target genes (red color-bar), cell cycle (blue color-bar) and immune function (green color-bar) with the indicated treatments across cell line models. **F.** Heatmap showing the log fold change of representative MEK target genes (red color-bar), cell cycle (blue color-bar) and immune function (green color-bar) with the indicated treatments across PDX models. **G.** Heatmap showing K-means clustering of TCGA pancreatic cancer cohort based on significant genes altered with palbociclib+trametinib related to immune function (red color-bar) and cell cycle (blue color-bar). **H.** Pearson correlation between Cluster 1 and Cluster 2 with a negative correlation of -0.48 and significance p<2.2e-16.(Right) **I.** Kaplan-Meier analysis for overall survival comparing Cluster 1 and Cluster 2 from the heatmap.

To functionally assess how the combination of CDK4/6 and MEK inhibition impacts on the immune system, we employed a syngeneic pancreatic cancer model derived from the KPC mouse model (the 4662 model)^29^. This model is highly aggressive and neither CDK4/6 inhibition nor MEK inhibition alone had a significant impact on either cell growth or tumor growth in C57BL6/J mice (Fig 4A and 4B). However, consistent with the findings in the patient-derived models the combination treatment with MEK and CDK4/6 inhibition delayed the progression of this model, albeit even within 14 days there was tumor growth on treatment (Fig 4B). To define how CDK4/6 and MEK inhibition impacts on the immunological tumor microenvironment and immune-checkpoint inhibitor therapy single cell sequencing was performed. Mice bearing established 4662 tumors were treated with palbociclib and trametinib, or a triple combination with anti-PD-L1. Tumors were dissociated and the CD45+ fraction was isolated by fluorescent-activated cell sorting (Fig 4C). Single cell RNA sequencing was performed using the 10x chromium method. Consistent with the enrichment approach, all cells were positive for the CD45 gene (PTPRC) irrespective of treatment (Fig 4C). Dimensional clustering algorithms defined 12 populations of cells that capture lymphoid, myeloid, monocytic, and B-cell components of the tumor (Fig 4D and S9). The myeloid compartment was defined by classical phenotyping markers (CD33, S100A8, and ITGAM), with two dominant myeloid-macrophage populations (clusters 0 and 2) (Fig 4E). Treatment with palbociclib and trametinib lead to a complete switch in the dominant macrophage population present in the control tumors (Fig 4F and G). Specifically, the myeloid-macrophages were suppressed for immediate early genes (e.g. EGR1, JUNB, ETS2, FOS) that are downstream from MEK/ERK signaling^44^. These changes were accompanied by down-regulation of PTGS2, VEGFA, and MMP9 that are associated with immune-suppressive M2-like macrophages (Fig 4H and S10)^45^. Concordantly, with treatment there was induction of genes associated with iron-metabolism (FTL1, FLH1, and HMOX1) and macrophage functions (BNIP3L, CTSD) that are typically associated with immune-activating M1-like macrophages (Fig 4H)^45^. Additionally, top induced genes include CCL3, CCL4, and CCL6 that are involved in T-cell, dendritic cell, B-cell, and NK cell infiltration (Fig 4H). This shift in macrophage populations was also observed in tumors treated with the triplet that includes anti-PD-L1 (Fig 4G and H). Under these treatments there was only a modest alteration in the neutrophil population (cluster 3).

**Figure 4.**
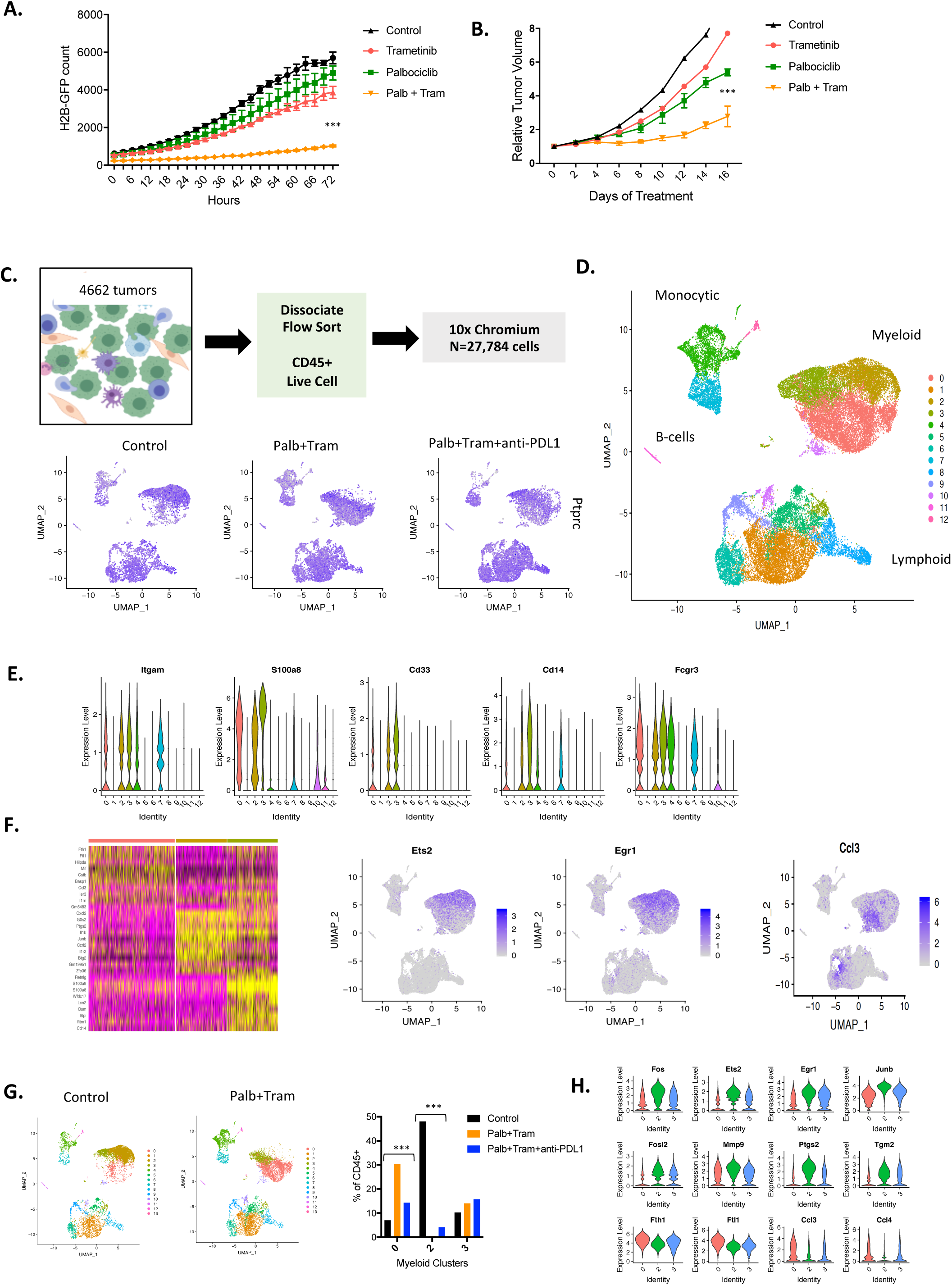
Impact of CDK4/6 and MEK inhibition on the tumor microenvironment: **A.** The KPC-derived 4662 cell line labeled with H2B-GFP was treated with palbociclib, trametinib or the combination. Cellular proliferation was determined using IncuCyte live cell imaging over the indicated time frame (***p<0.001). **B.** The 4662 model was introduced into C57/BL6J mice subcutaneously, mice were randomized at approximately 200 mm^3^ and tumor volume was measured by calipers (***p<0.001). **C.** Schematic workflow for the single cell sequencing. Feature plots denote the expression of Ptprc (CD45 gene) in all of the captured cells with treatment. **D.** All of the cells from the treated mice were filtered for quality and clustered using Seurat software in the dimension plot. The general clustering of different immunological subtypes is indicated. **E.** The violin plots denote gene markers associated with myeloid, neutrophil, and macrophage populations. **F.** Seurat heatmap identifying top differences between the myeloid clusters and representative feature plots are shown. **G.** Distribution of cells in the indicated myeloid clusters, changes in clusters 0 and 2 with treatments are highly significant by chi-square statistic (***p<1E-100). **H.** Genes differentially expressed in cells between clusters 2 and 0 are highlighted in the violin plot (all genes are highly significant between the clusters p<1E-5).

The lymphoid compartment of the tumor encompassed multiple sub-types of T-cells and NK cells as indicated by conventional markers (Fig 5A, 5B, S11). With treatment the lymphoid populations were clearly enhanced with both CDK4/6 and MEK inhibition as well as the triplet combining these inhibitors with anti-PD-L1 (Fig 5C). The most common CD8+ T-cell population (cluster 1) almost doubled with treatment of palbociclib and trametinib and was further enhanced with anti-PD-L1. This T-cell population is GZMB and PRF1 positive, characteristics of cytotoxic function. Interestingly, other populations were more modestly induced or did not change with the doublet (Fig 5D). This includes NK-cells and a proliferative T-cell population (cluster 9) which was surprisingly unaffected by CDK4/6 and MEK inhibition (Fig 5D and S12). The inclusion of anti-PD-L1 treatment further enhanced the T-cell infiltrate with significant increases in veritably all populations, most notably the NK cell population (Fig 5D). In the control tumors, there was a significant skewing toward myeloid infiltration, while treatment with CDK4/6 and MEK inhibitors increases the lymphoid infiltrate which, was further enhanced with anti-PDL1 (Fig 5E). These findings were confirmed in independent tissues using anti-CD8 and CD163 immunostaining and multi-spectral imaging (Fig 5F and S12). The monocytic populations were more limited in the treatment-naïve tumors, but augmented with treatment, as were relatively rare BATF3+ dendritic cell populations (Fig S13), Velocity analysis suggests that the monocytic pools were transiting toward the dendritic cell populations (Fig S13). In addition, while limited, the number of B-cells within the tumor increased. Together, these data suggest an overall increase in tumor immunogenicity with CDK4/6 and MEK inhibition. The 4662 syngeneic model is largely resistant to anti-PD-L1 treatment consistent with other studies^29^; however, the combination of CDK4/6 and MEK inhibition with anti-PD-L1 was highly effective yielding regression of the tumor (Fig 5G). This treatment was well-tolerated as indicated by lack of weight loss (Fig S14), and the combination of both CDK4/6 and MEK inhibitor was required for the potent cooperation with anti-PD-L1 (Fig S14). In a small cohort of animals followed long-term, tumor cure and resistance to subsequent challenge was also observed (Fig S14). To confirm the activity, we employed orthotopic models, where tumor cells were introduced into the pancreas and tumor engraftment and volume was assessed by magnetic resonance imaging (MRI) (Fig. 5H). Mice were randomized to treatment groups, and as in the subcutaneous models, there was more profound disease control with the triplet combining CDK4/6 and MEK inhibitors and anti-PD-L1. To specifically interrogate the significance of the CD8+ T-cells for this therapeutic response, mice were coordinately treated with anti-CD8, which depleted CD8+ cells in the tumor bed (Fig S14), and reversed the impact of anti-PD-L1 (Fig 5H). Together, these studies underscore the potency of combining targeted and immunotherapy for the treatment of PDAC and illuminate previously unrecognized contributions of targeted therapies on the tumor-immune microenvironment.

**Figure 5.**
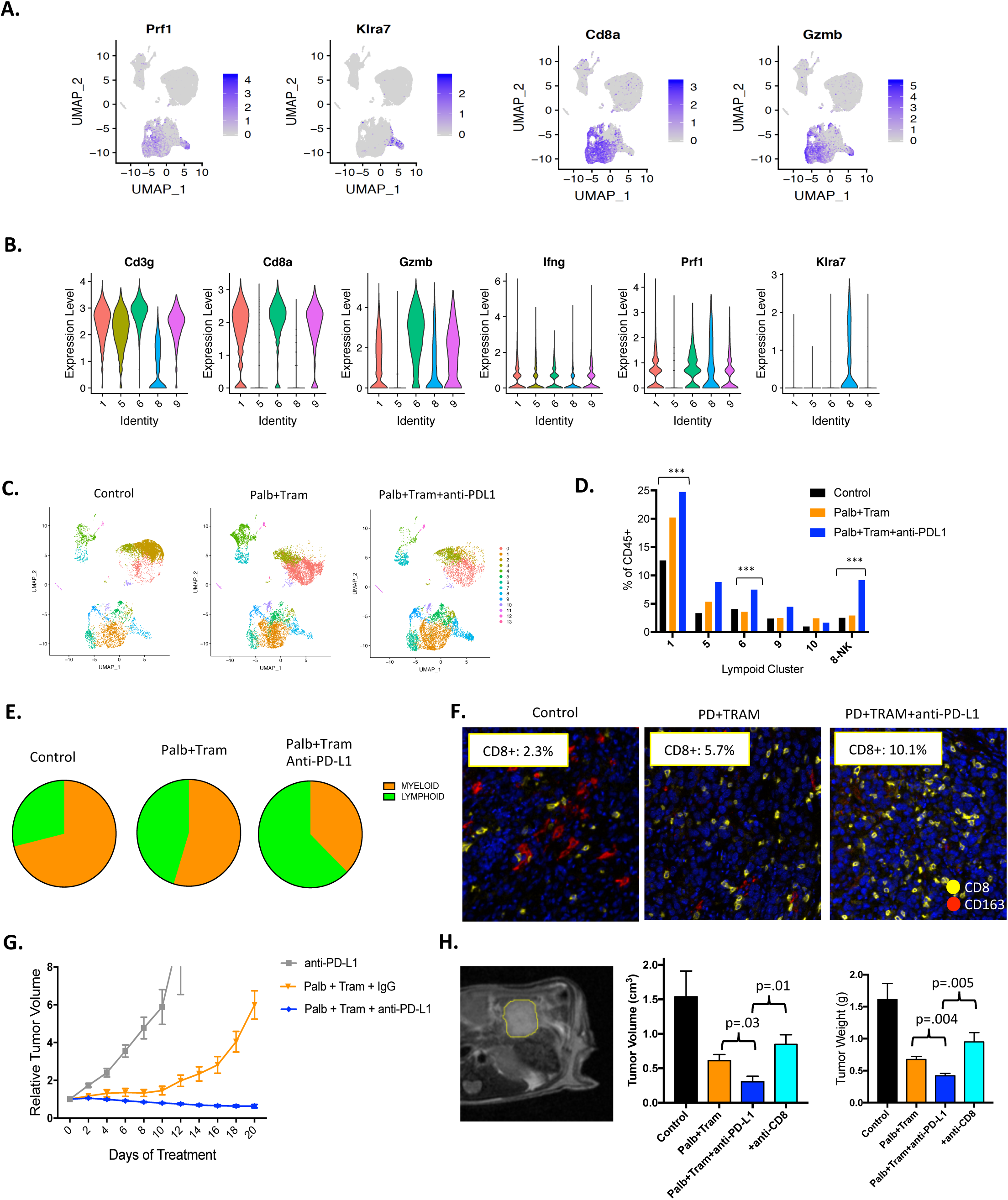
Cooperation between CDK4/6-MEK inhibition and immune-checkpoint inhibition: **A.** Predominant T-cell clusters were defined using feature plots of classical markers. **B.** Violin plots demonstrating the cellular expression of the indicated markers across the predominant lymphoid clusters. **C.** Dimension cluster plots illustrating the accumulation of T-cell populations with the indicated treatments. **D.** Quantitation of cells present in each of the T-cell and NK cell clusters is shown. Changes indicated are statistically significant as determined by Chi-squared analysis (p<1E-100). **E.** A proliferative CD8+ T-cell population is represented in cluster 9, the expression of Ki67 in CD8+ T-cells was confirmed by multi-spectral imaging. **F.** Myloid to lymphoid relationship as determined by single cell sequencing analysis **G.** Tumor volume was determined in the 4662 model treated with the indicated regimens (***p<0.001). **H.** Representative axial T2-weighter magnetic resonance image showing orthotopically engrafted 4662 tumor model. Mice were treated as indicated and anti-CD8 was employed to neutralize the T-cell response in the triplet combination.

## DISCUSSION

As a therapy recalcitrant disease, combinatorial means to target pancreatic cancer will assuredly be required. While CDK4/6 inhibitors have been found to be effective in ER+ breast cancer and FDA-approved for that indication, clinical advances in additional indications have yet to mature^46^. As a tumor driven by KRAS and CDKN2A loss, it would be expected that CDK4/6-inhibition would have dominant effects in pancreatic cancer. The data herein reinforce the finding that cell cycle plasticity enables escape from CDK4/6 inhibitors; however, by unbiased drug screens MEK inhibitors emerged as key cooperative agents to block that plasticity and couple CDK4/6 inhibition to the activation of RB, suppression of CDK2, and repression of E2F target genes. These interactions are synergistic in cell culture and highly potent in all PDX models representing 10 different patients. While the combination is highly effective, with cessation of treatment cells will re-enter the cell cycle (not shown) and tumors ultimately progress. Additionally, in very fast growing tumors e.g. 4662 model, progression occurs on therapy and reinforces the need for an adjuvant strategies to complement the cell cycle inhibition.

In evaluating the effect of CDK4/6 and MEK inhibition on gene expression, we observed a large number of genes that are repressed or activated and could yield a specific vulnerability beyond the cell cycle. For example, there is potent suppression of EZH2 and additional chromatin modifiers as well as inhibition of DNA repair mechanisms that could confer sensitivity to sensitivity to diverse therapeutics. A noted feature of the response to CDK4/6 and MEK inhibition is the upregulation of interferon and antigen presentation that has been observed in other tumor types^27, 47^. This signature is related to cell cycle arrest and is RB-dependent; however, whether it is related to senescence is unclear in this setting. The gene expression analysis of cell lines occurred at 48-72 hours before the morphological changes associated with senescence, additionally the signature defined here shares little similarity with the classical SASP signature defined by Dr. Campisi and colleagues^26, 48^. Additionally, while the combination of CDK4/6 and MTOR inhibitors induces potent cell cycle arrest, the induction of immune-related pathways is not present. We favor the model that aberrant chromatin/transcripts elicited as a consequence of RB activation in the presence of oncogenic signals drive the interferon-like response, which is highly related to that mediated by knockdown of EZH2 or loss of DNMT1 which are both repressed with MEK and CDK4/6 inhibition^26, 49^.

The impact of targeted agents on the immune system is becoming progressively important with the success of immune-checkpoint therapies. Since both MEK and CDK4/6 have important roles in hematopoiesis understanding the intersection with the immunological milieu within the tumor is clearly significant^50^. Using single cell sequencing of CD45+ cells provided a highly detailed analysis of the immune repertoire in a KPC-based tumor model. These tumors are dominated by myeloid populations, which is consistent with data mouse models and clinical pancreatic cancer cases^24, 51^. Treatment with CDK4/6 and MEK inhibition had a pronounced effect on the immune cells within the tumor with the most notable changes occurring within the myeloid-macrophage population. In this context, the combinatorial treatment limited immediate early genes that are associated with angiogenesis and pro-tumorigenic elements of M2-macrophages. These changes were linked to the acquisition of a macrophage population that exhibited enhanced iron-metabolism and chemokine secretion promoting recruitment of lymphocytes and antigen presenting cells. These proximal alterations on immediate early signaling, were associated with increased recruitment of selected T-cell populations. However, for potent engagement of NK populations into the tumor anti-PDL1 was required. These data are consistent with the cooperative effect of CDK4/6-MEK inhibition with anti-PDL1. This cooperation was T-cell mediated as anti-CD8 treatment limits the effectiveness of therapy. Furthermore, in mice that were cured by the triplet therapy there was subsequent anti-tumor immunity. These data suggest that the targeted therapies utilized herein can serve as a significant adjunct to immune-checkpoint inhibition in RAS-driven tumors.

## Notes

CONFLICT OF INTEREST: None of the authors have a conflict of interest to report

