## Supplementary Figures for "Targeting dual signaling pathways in concert with immune checkpoints for the treatment of pancreatic cancer"

**Supplemental Figure Legends:**

**Figure S1. A.** The 3226 cell line was treated with DMSO or Palbociclib (100nM) for 24 hours before addition of drug library consisting of over 300 cancer drugs at 100nM. After 72 hours, relative viability was calculated by H2B-GFP analysis. Graph shows relative cell growth of agent alone with DMSO (black) or agent with Palbociclib (grey). MEK inhibitors with Palbociclib are highlighted in orange. **B.** Isobologram analysis of palbociclib and trametinib dose responses, in all cell models the drug interaction was synergistic for the suppression of BrdU incorporation. **C.** Immunoblot analysis to determine the effect of palbociclib (100 nM) +/- trametinib (100 nM) on the mitogenic signaling pathways from the indicated cell lines following 48 h exposure.

**Figure S2. A.** Fluorescence ubiquitination cell cycle indicator (FUCCI) labeled 226 cells were treated with DMSO, Palbociclib 200nM (Palb), Trametinib 50nM (Tram), and combination (Palb + Tram) for 48 hours. Panels show GFP on y-axis and RFP on x-axis. Lower panel shows DNA content as determined by DAPI counter staining. **B.** Immunoblot analysis of p27 expression with increasing doses of trametinib and the combination as indicated. **C.** CDK2 kinase assays against an exogenous RB substrate are shown. The quantitation is from two independent experiments (*$p<0.05$).

**Figure S3.** Representative immunohistochemical staining for phospho-RB, Cyclin D1, and Cyclin E. Two independent PDX models treated with palbociclib or the combination with trametinib are shown.

**Figure S4. A.** Heatmaps showing the logFC of genes that are reproducibly repressed and induced with palbociclib and trametinib treatments in two different cell lines. The columns show independent RNA sequencing samples. **B.** Barplots show the changes of gene expression of select genes in different cell models with the indicated drug treatments. The 7310 cell line is RB-deficient and serves as a negative control. **C.** Representative immunofluorescence staining for HLA-A and HLA-C in 519 and 7310 cell models. The 7310 cell line is RB-deficient and serves as a negative control.

**Figure S5. A.** The secretion of CCL5 and CXCL10 was determined by ELISA assay from media from the indicated cells. These two chemokines were significantly induced by treatment with palbociclib and trametinib (***$p<0.001$). **B.** Representative gene set enrichments between the cell line and PDX from the same model are shown. **C.** Volcano plots show the behavior of consensus palbociclib and trametinib repressed (blue) and induced (green) gene defined in cell lines, in the context of all the PDX models utilized.

**Figure S6.** The volcano plots show the behavior of the canonical SASP signature from Coppe et al. in PDX models treated with palbociclib (green) or palbociclib and trametinib (orange).

**Figure S7. A.** Heatmaps from the indicated cell lines and PDX models treated with palbociclib (green colorbar) or the combination with trametinib (orange colorbar) or TAK228 (purple colorbar). **B.** The logFC of select genes under the indicated conditions.

**Figure S8.** Heatmaps show the behavior of immune or cell cycle genes in the context of the TCGA pancreatic cancer gene expression data, related Kaplan-Meier plots are shown.

**Figure S9.** Seurat heatmap showing the top differentially expressed genes in the 12 clusters from the single cell sequencing data from N=27,784 cells.

**Figure S10.** **A.** Feature plots focused on genes that are distinct between myeloid clusters in control and palbociclib+trametinib treated tumors. **B.** Violin plots of the indicated genes across all clusters.

**Figure S11.** Violin plots of genes associated with T-cell function across all of the clusters.

**Figure S12.** **A.** Representative images of CD8 staining with the indicated treatments in treated tumor tissue. **B.** Fraction of CD163 to CD8 staining determined using Aperio image analysis of 4662 tumors. **C.** Cluster 9 is the only immunological cell type that is proliferative. This cluster remains present with CDK4/6 and MEK inhibitor treatment and is enhanced with anti-PD-L1. Representative images of Ki67 staining in CD8+ T-cells.

**Figure S13.** **A.** The percentage of cells in the monocytic clusters 4 and 7 are shown. Violin plots show gene expression related to HLA and antigen presentation. **B.** Velocity analysis illustrating that these monocytic populations are differentiating toward dendritic cell population **C.** The percentage of cells in the B-cell and dendritic cell clusters is shown. Violin plots showing B-cell and dendritic cell markers across all clusters.

**Figure S14.** **A.** Mice were randomized when tumor volumes reached $150mm^3$ and treated with vehicle (control), Palbociclib and Trametinib, or the combination with and anti-PD-L1 for 21 days. Tumor weight at sacrifice is shown (***$p<0.001$). **B.** Body weight of mice in indicated treatments of vehicle (control), Palbociclib and Trametinib, and combination with anti-PD-L1. **C.** Mice were randomized when tumor volumes reached $150mm^3$ and treated with vehicle (control), Trametinib and anti-PD-L1 (Tram + anti-PD-L1), or Palbociclib and anti-PD-L1 (Palb + anti-PD-L1) for 21 days. **D.** Analysis of CD8+ cells within the tumor bed by Immunohistochemistry. Cells were quantified by Aperio image analysis. **E.** Tumor volume from mice treated with vehicle or completely regressed on treatment with palbociclib and trametinib and anti-PD-L1. Mice were re-challenged with tumor inoculation as shown.

Figure S1

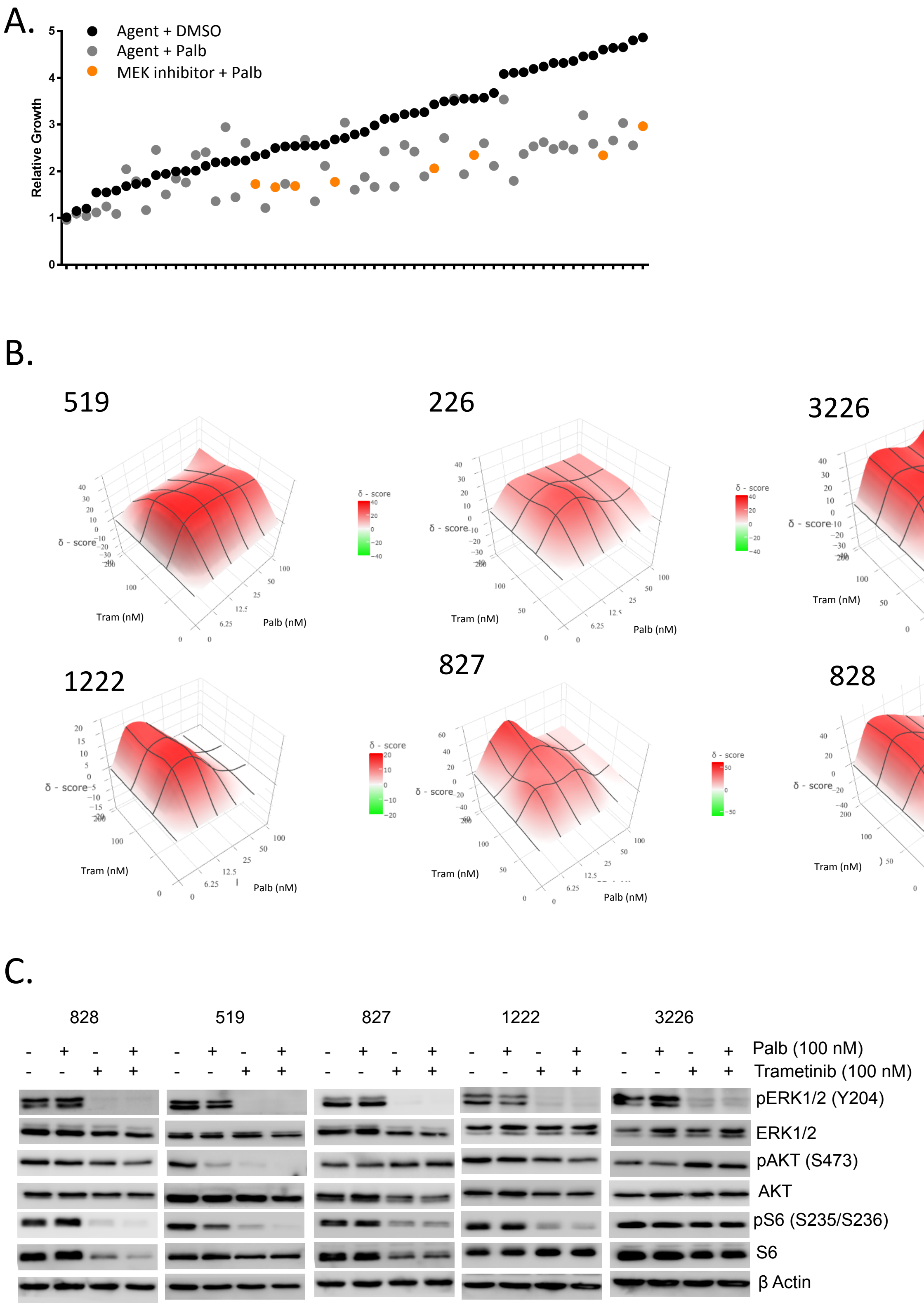
A.
Agent + DMSO
Agent + Palb
MEK inhibitor + Palb
Relative Growth
B.
519
226
3226
1222
827
828
δ - score
Tram (nM)
Palb (nM)
C.
828
519
827
1222
3226
Palb (100 nM)
Trametinib (100 nM)
pERK1/2 (Y204)
ERK1/2
pAKT (S473)
AKT
pS6 (S235/S236)
S6
β Actin

Figure S2

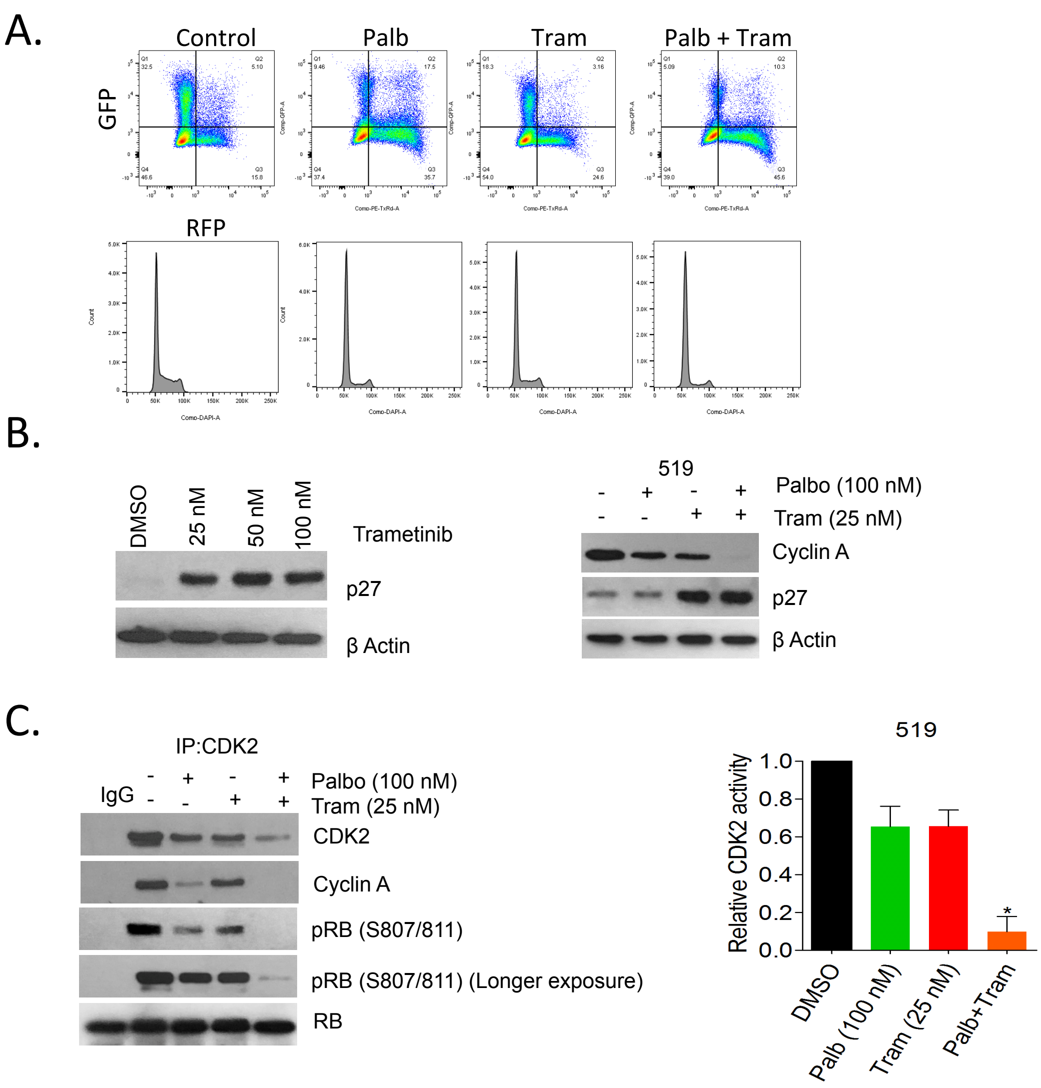

A.
Control
Palb
Tram
Palb + Tram
GFP
RFP
B.
DMSO
25 nM
50 nM
100 nM
Trametinib
p27
β Actin
519
Palbo (100 nM)
Tram (25 nM)
Cyclin A
p27
β Actin
C.
IP:CDK2
IgG
Palbo (100 nM)
Tram (25 nM)
CDK2
Cyclin A
pRB (S807/811)
pRB (S807/811) (Longer exposure)
RB
519
Relative CDK2 activity
1.0
0.8
0.6
0.4
0.2
0.0
DMSO
Palb (100 nM)
Tram (25 nM)
Palb+Tram

Figure S3

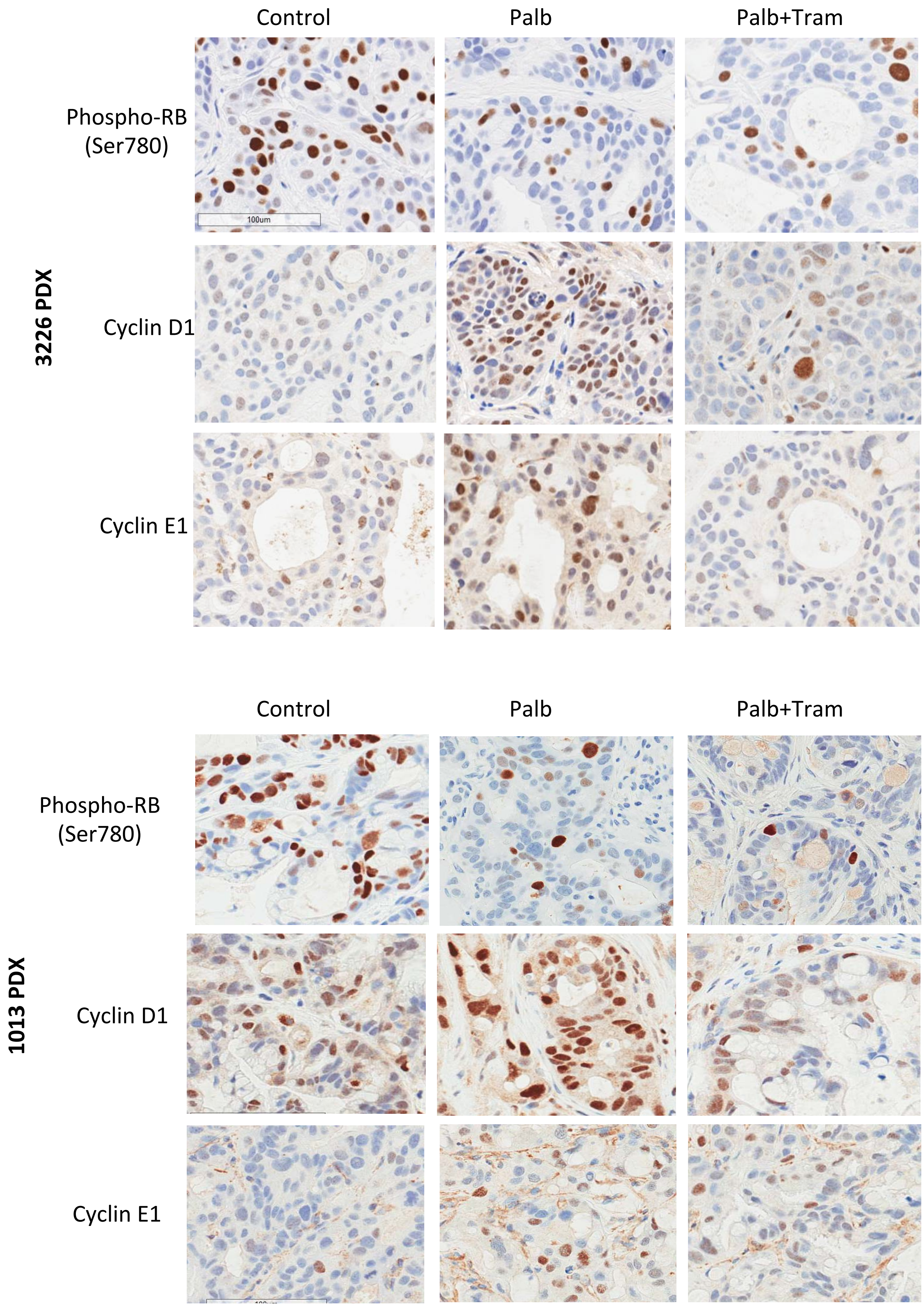

Control
Palb
Palb+Tram
3226 PDX
Phospho-RB (Ser780)
100um
Cyclin D1
Cyclin E1
Control
Palb
Palb+Tram
1013 PDX
Phospho-RB (Ser780)
Cyclin D1
Cyclin E1

Figure S4

A.

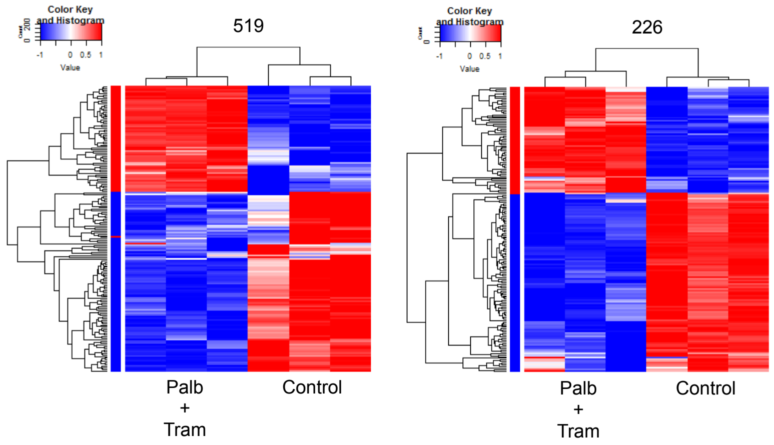


B.

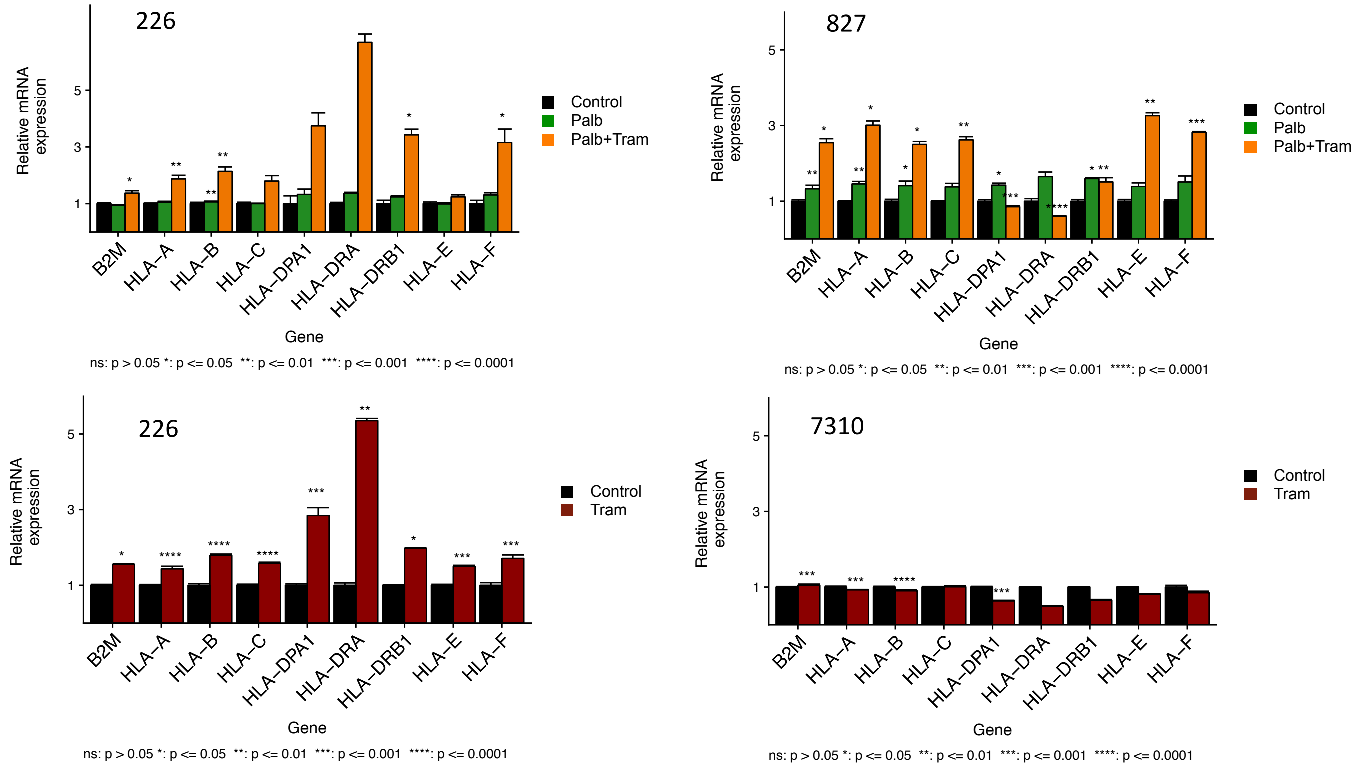


C.

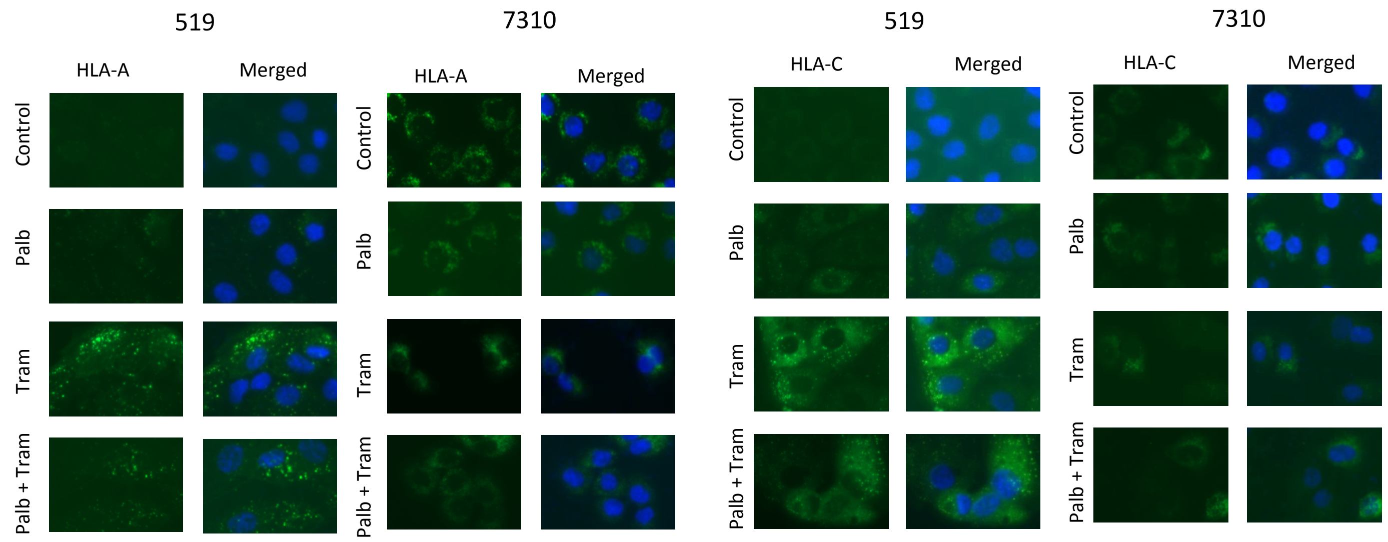

Figure S5

A.

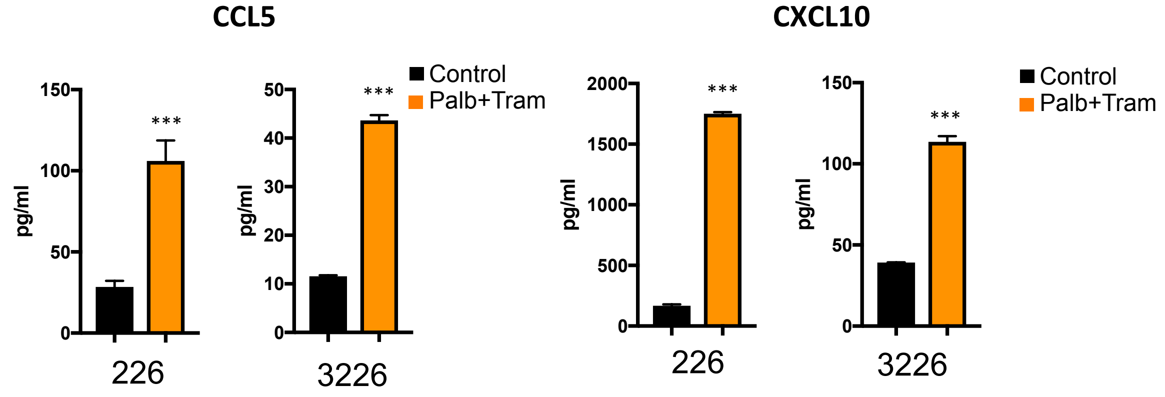


B.

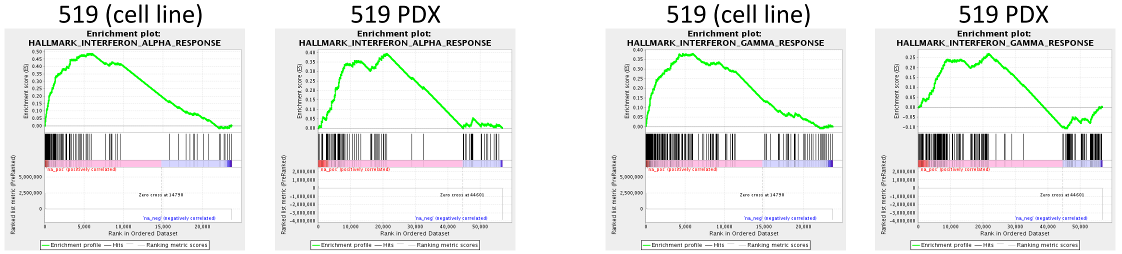


C.

PDX cohort (Palb+Tram): cell culture signature

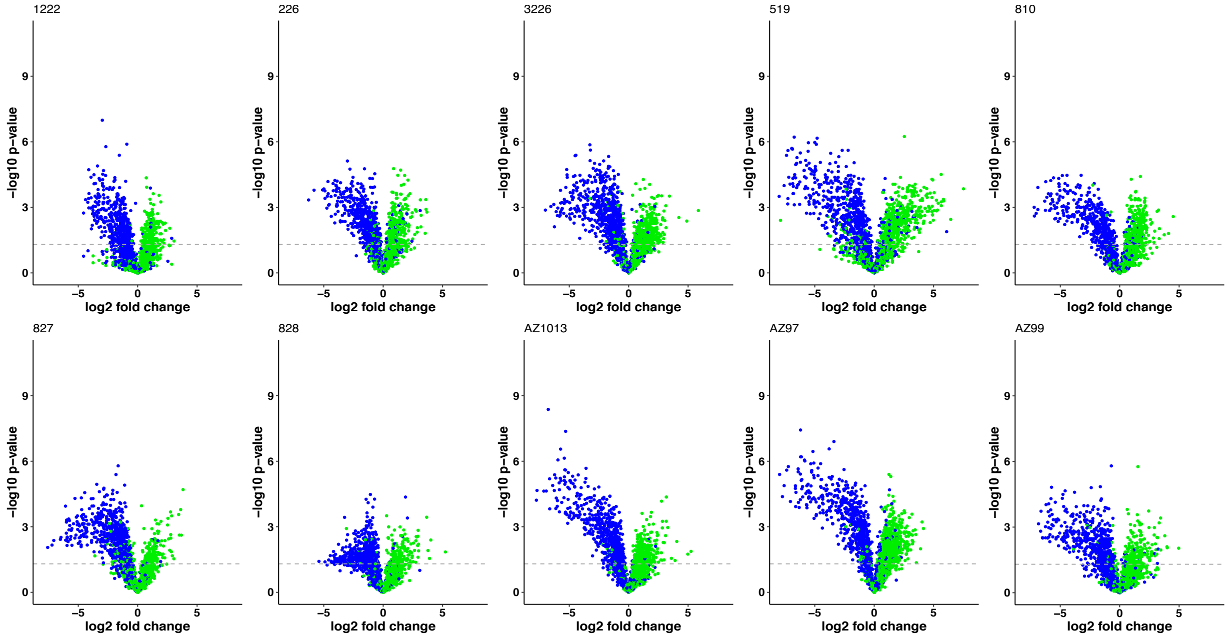

### PDX cohort (Palb): SASP Signature

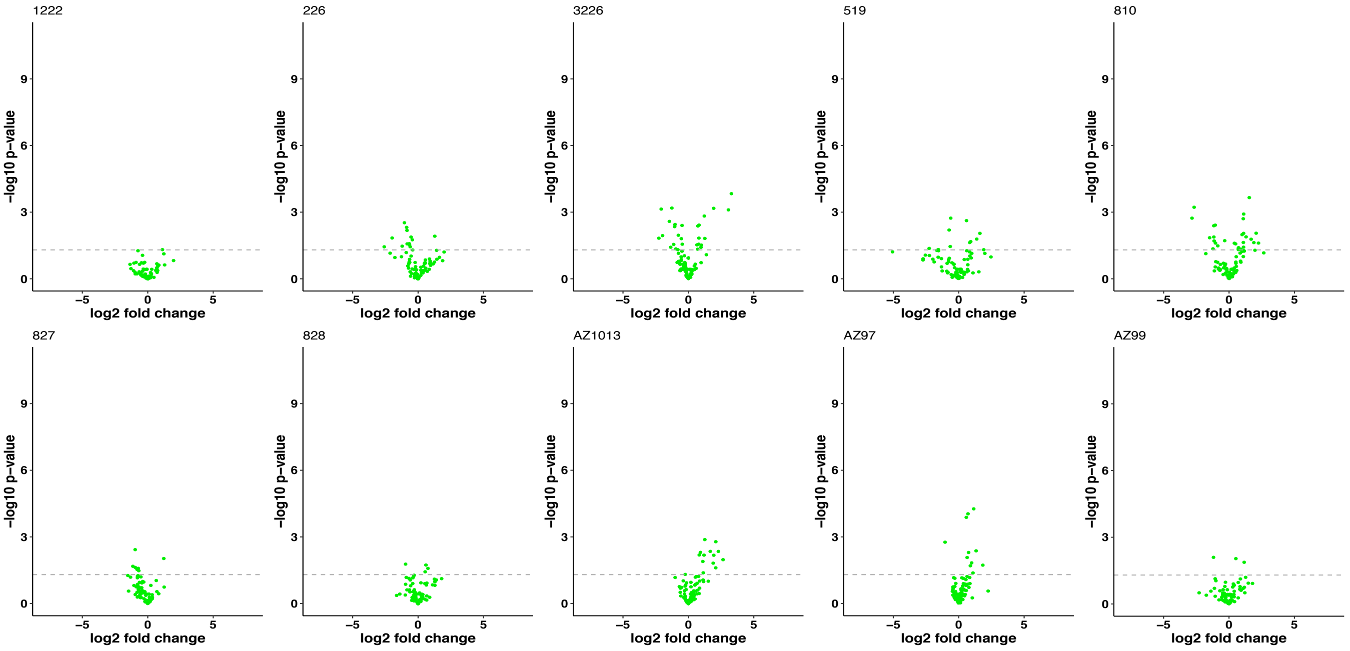


### PDX cohort (Palb+Tram): SASP Signature

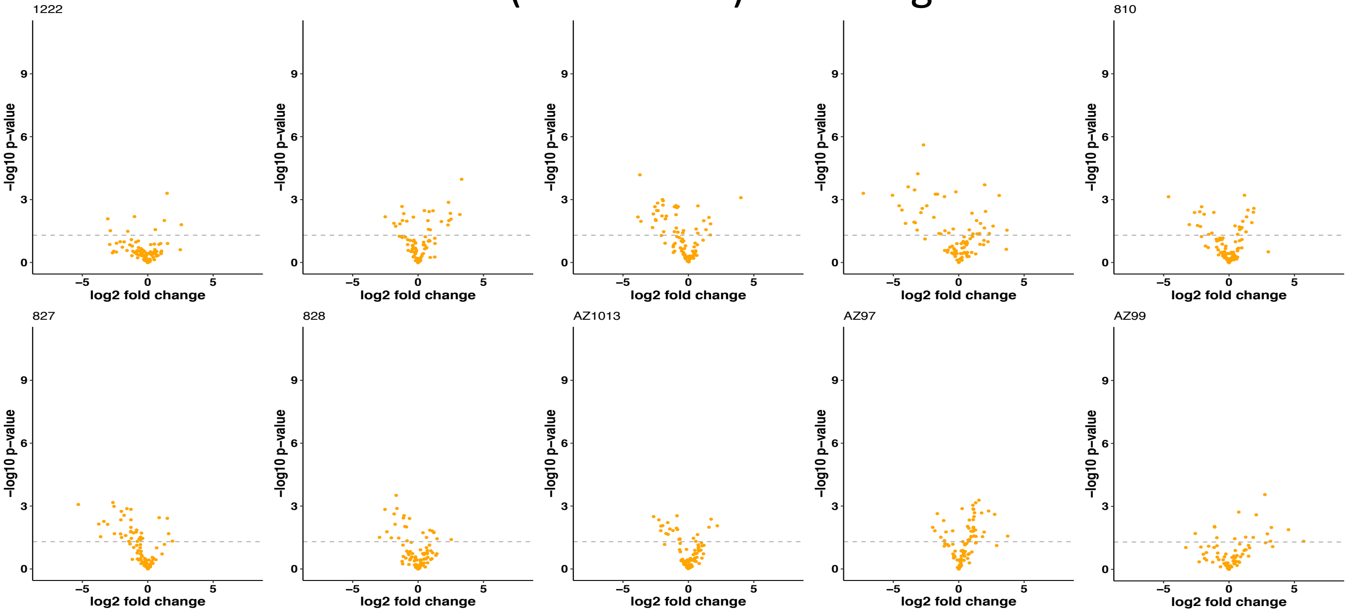

A.

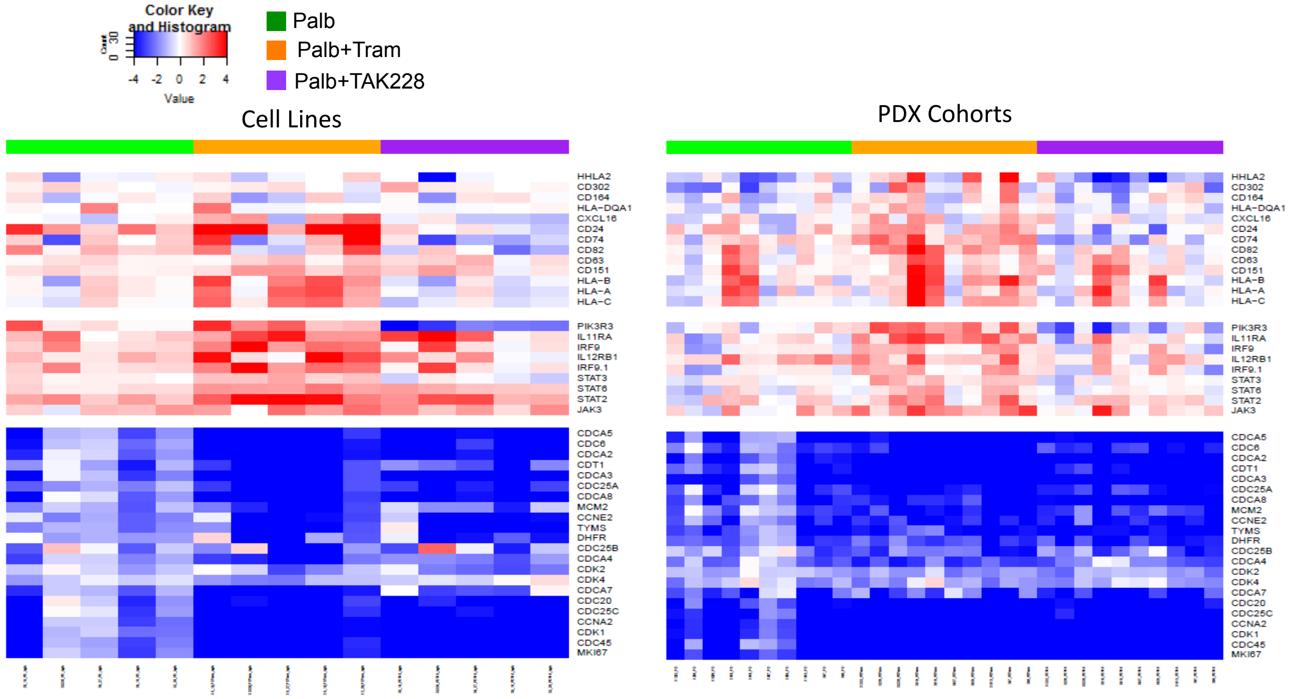
Color Key and Histogram
Count
Value
Palb
Palb+Tram
Palb+TAK228
Cell Lines
PDX Cohorts
HHLA2
CD302
CD164
HLA-DQA1
CXCL16
CD24
CD74
CD82
CD63
CD151
HLA-B
HLA-A
HLA-C
PIK3R3
IL11RA
IRF9
IL12RB1
IRF9.1
STAT3
STAT6
STAT2
JAK3
CDCA5
CDC6
CDCA2
CDT1
CDCA3
CDC25A
CDCA8
MCM2
CCNE2
TYMS
DHFR
CDC25B
CDCA4
CDK2
CDK4
CDCA7
CDC20
CDC25C
CCNA2
CDK1
CDC45
MKI67

B.

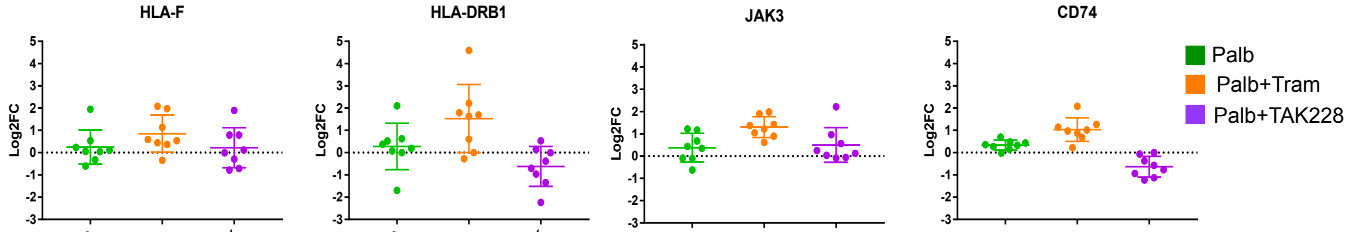
HLA-F
HLA-DRB1
JAK3
CD74
Log2FC
Palb
Palb+Tram
Palb+TAK228

Figure S8

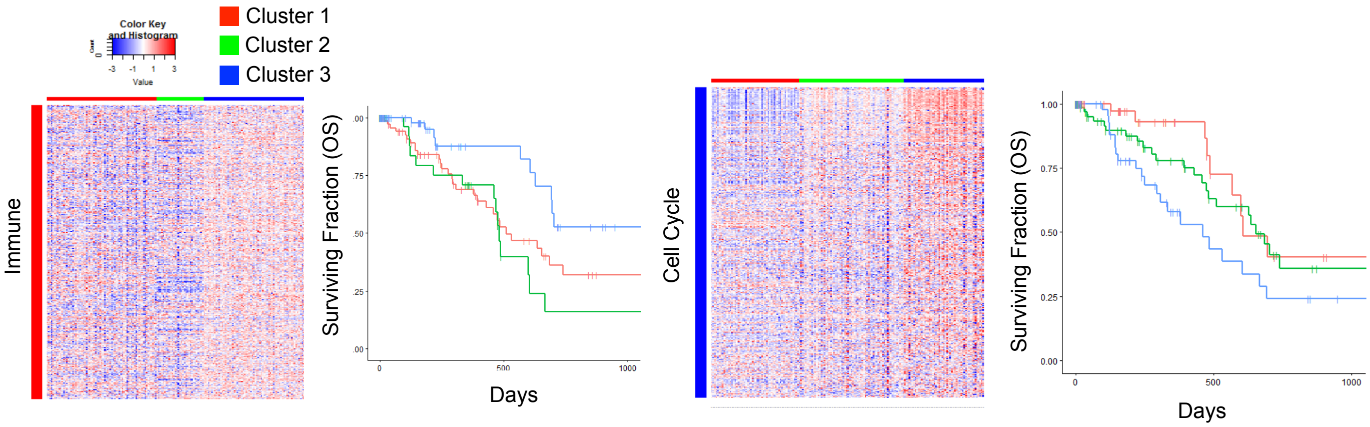

Color Key and Histogram
Count
Value
Cluster 1
Cluster 2
Cluster 3
Immune
Surviving Fraction (OS)
Days
Cell Cycle
Surviving Fraction (OS)
Days

Figure S9

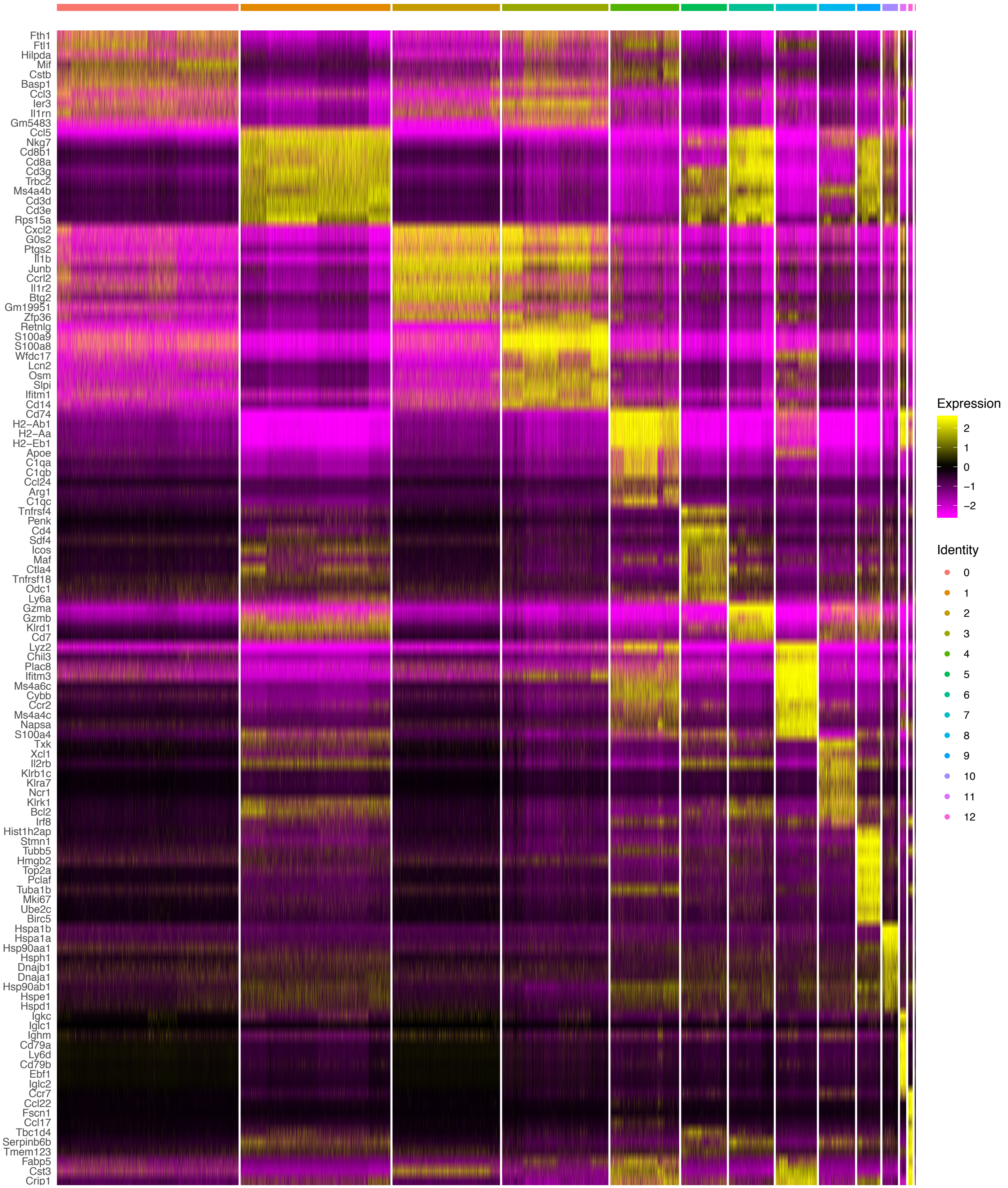
Fth1
Ftl1
Hilpda
Mif
Cstb
Basp1
Ccl3
Ier3
Il1rn
Gm5483
Ccl5
Nkg7
Cd8b1
Cd8a
Cd3g
Trbc2
Ms4a4b
Cd3d
Cd3e
Rps15a
Cxcl2
G0s2
Ptgs2
Il1b
Junb
Ccrl2
Il1r2
Btg2
Gm19951
Zfp36
Retnlg
S100a9
S100a8
Wfdc17
Lcn2
Osm
Slpi
Ifitm1
Cd14
Cd74
H2−Ab1
H2−Aa
H2−Eb1
Apoe
C1qa
C1qb
Ccl24
Arg1
C1qc
Tnfrsf4
Penk
Cd4
Sdf4
Icos
Maf
Ctla4
Tnfrsf18
Odc1
Ly6a
Gzma
Gzmb
Klrd1
Cd7
Lyz2
Chil3
Plac8
Ifitm3
Ms4a6c
Cybb
Ccr2
Ms4a4c
Napsa
S100a4
Txk
Xcl1
Il2rb
Klrb1c
Klra7
Ncr1
Klrk1
Bcl2
Irf8
Hist1h2ap
Stmn1
Tubb5
Hmgb2
Top2a
Pclaf
Tuba1b
Mki67
Ube2c
Birc5
Hspa1b
Hspa1a
Hsp90aa1
Hsph1
Dnajb1
Dnaja1
Hsp90ab1
Hspe1
Hspd1
Igkc
Iglc1
Ighm
Cd79a
Ly6d
Cd79b
Ebf1
Iglc2
Ccr7
Ccl22
Fscn1
Ccl17
Tbc1d4
Serpinb6b
Tmem123
Fabp5
Cst3
Crip1
Expression
2
1
0
−1
−2
Identity
0
1
2
3
4
5
6
7
8
9
10
11
12

A.

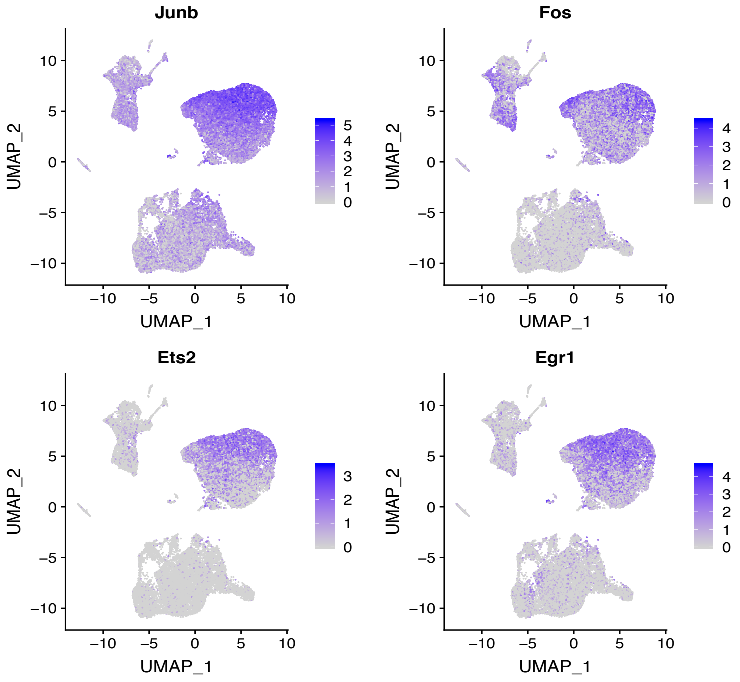
Junb
Fos
Ets2
Egr1
UMAP_1
UMAP_2

B.

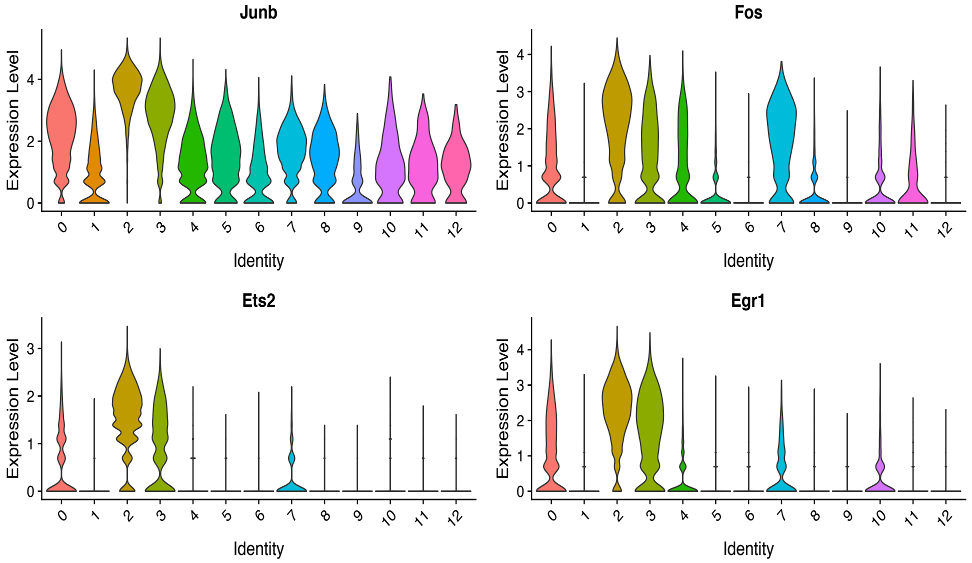
Junb
Fos
Ets2
Egr1
Expression Level
Identity

Figure S11

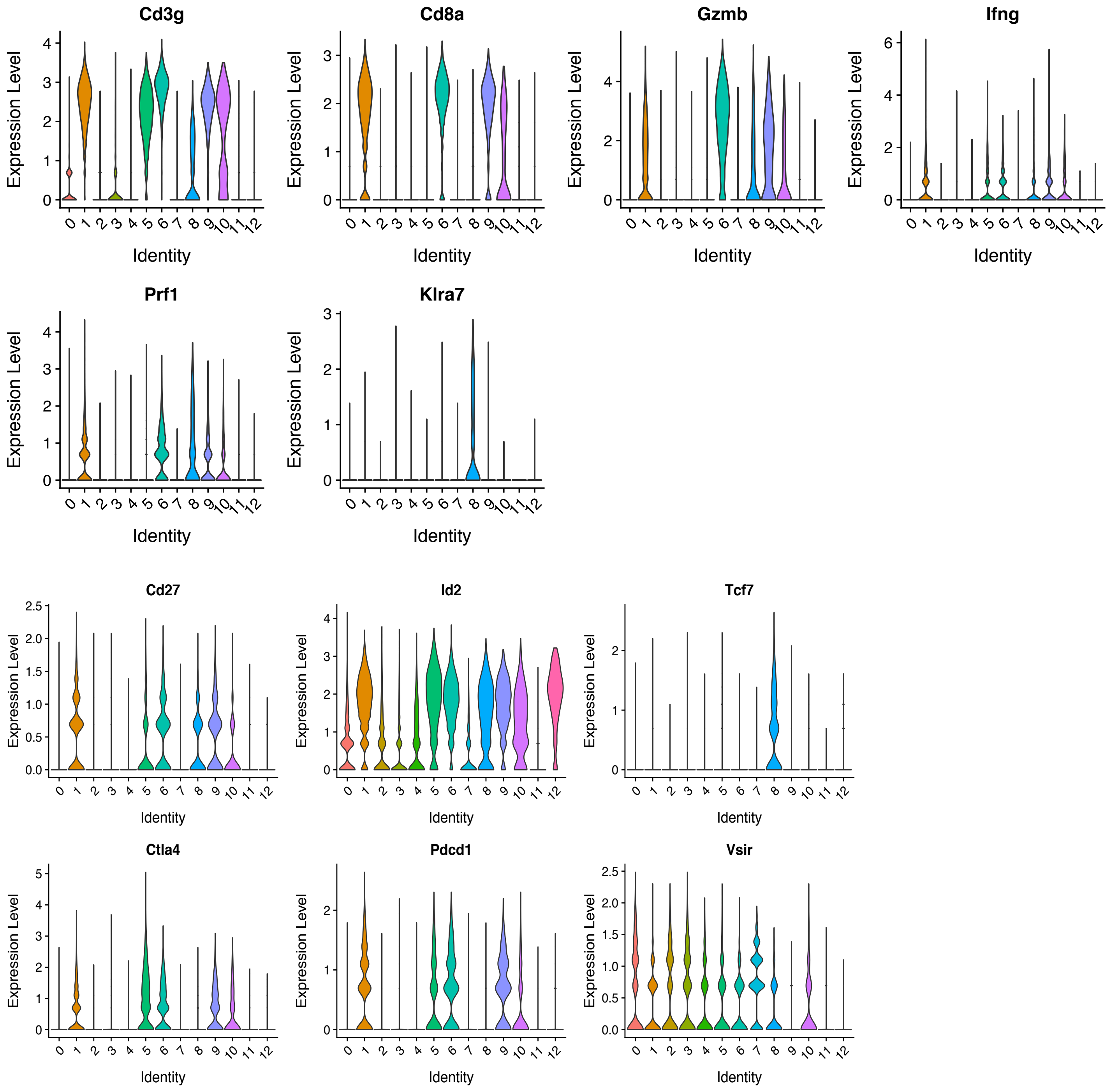

Cd3g
Cd8a
Gzmb
Ifng
Prf1
Klra7
Cd27
Id2
Tcf7
Ctla4
Pdcd1
Vsir
Expression Level
Identity
0 1 2 3 4 5 6 7 8 9 10 11 12

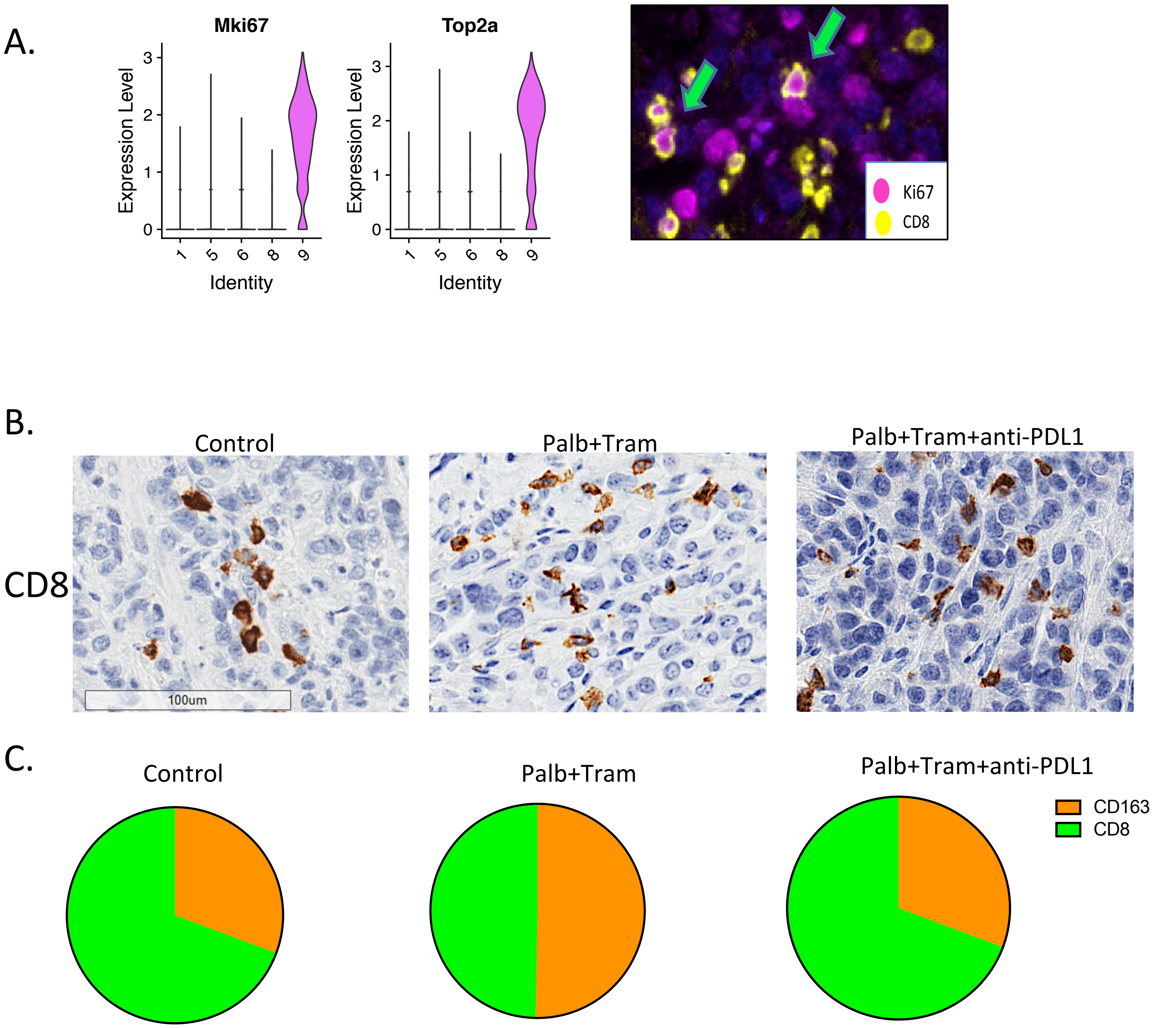
A.
Mki67
Top2a
Expression Level
Identity
1
5
6
8
9
Ki67
CD8
B.
Control
Palb+Tram
Palb+Tram+anti-PDL1
CD8
100um
C.
Control
Palb+Tram
Palb+Tram+anti-PDL1
CD163
CD8

Figure S13

A.

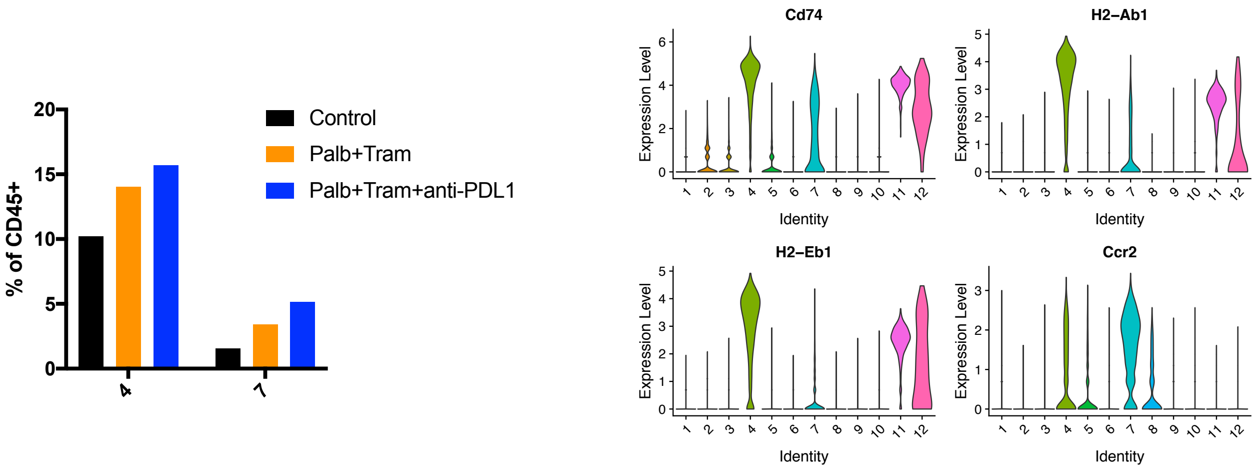


B.

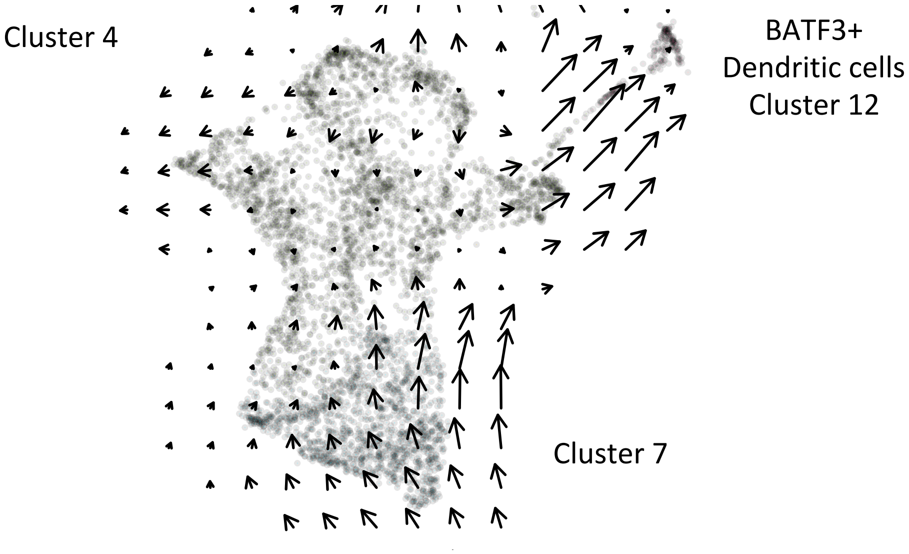


C.

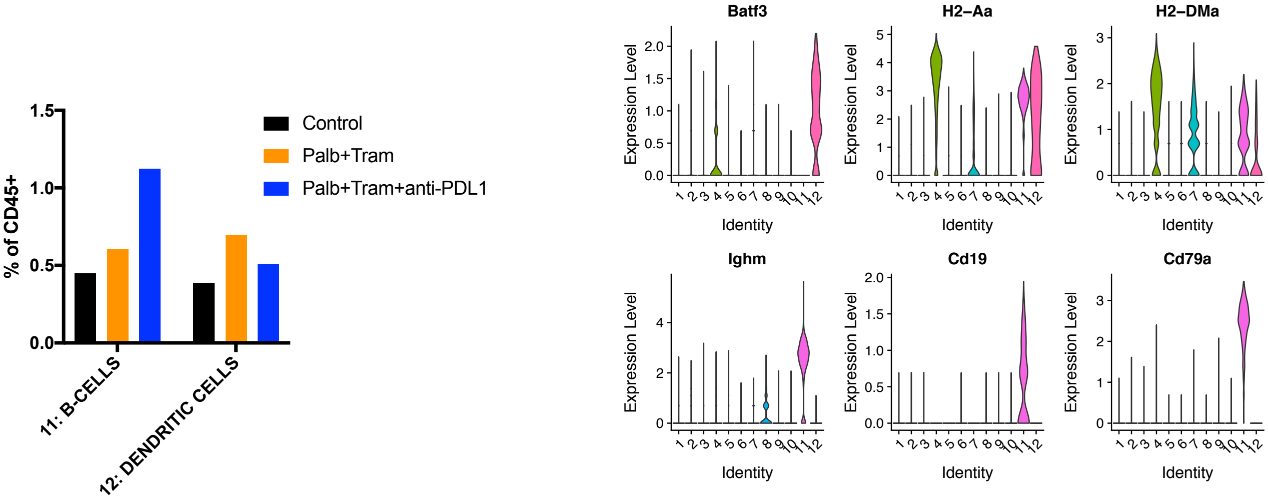

Figure S14

A.

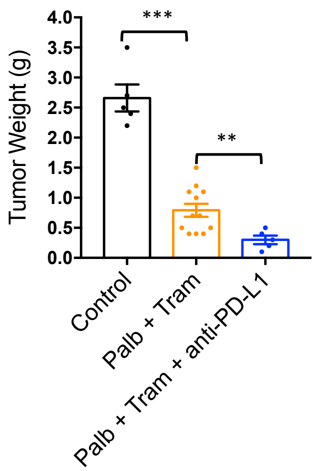


B.

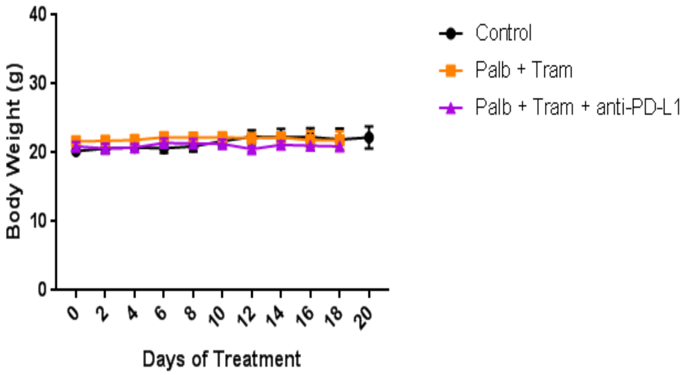


C.

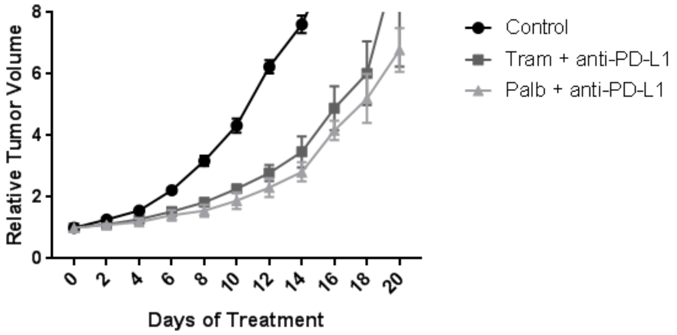


D.

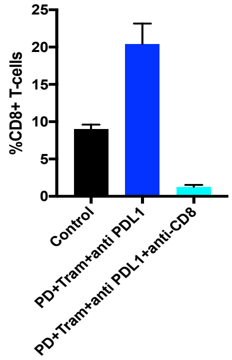


E.

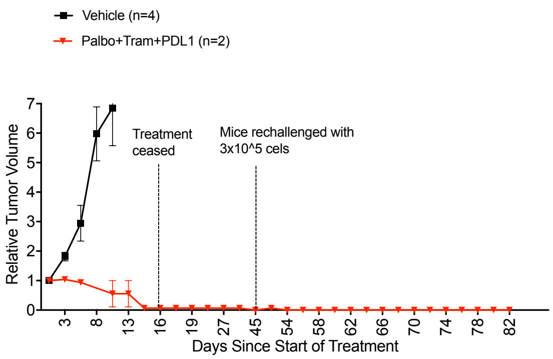
